# Amyloid fibril-based hydrogels for high-throughput tumor spheroid modeling

**DOI:** 10.1101/2020.12.28.424634

**Authors:** Namrata Singh, Komal Patel, Ambuja Navalkar, Pradeep Kadu, Debalina Datta, Debdeep Chatterjee, Abhishek Shaw, Sachin Jadhav, Samir K. Maji

## Abstract

Biomaterials mimicking extracellular matrices (ECM) for three-dimensional (3D) cultures have gained immense interest in tumor modeling and *in vitro* organ development. Here, we introduce versatile, thixotropic amyloid hydrogels as a bio-mimetic ECM scaffold for 3D cell culture as well as high-throughput tumor spheroid formation using a drop cast method. The unique cross-β-sheet structure, sticky surface, and thixotropicity of amyloid hydrogels allow robust cell adhesion, survival, proliferation, and migration, which are essential for 3D tumor modeling with various cancer cell types. The spheroids formed show overexpression of the signature cancer biomarkers and confer higher drug resistance compared to two-dimensional (2D) monolayer cultures. Using breast tumor tissue from mouse xenograft, we showed that these hydrogels support the formation of tumor spheroids with a well-defined necrotic core, cancer-associated gene expression, higher drug resistance, and tumor heterogeneity reminiscent of the original tumor. Altogether, we have developed a rapid and cost-effective platform for generating *in vitro* cancer models for the screening of anti-cancer therapeutics and developing personalized medicines.

## Introduction

Developing anti-cancer therapeutics generally uses a two-dimensional (2D) cell culture platform, which often provide ambiguous experimental observations due to different microenvironments experienced by cells in 2D versus three-dimensional (3D) tumor tissues^1–3^. Although cancer xenograft models using animals have great significance for the drug development process, the ethical problems, infrastructure, and extensive time requirement, limit their clinical usage^4^. To mimic the 3D microenvironment of solid tumors, previous studies have developed a 3D cell culture system, using biomaterials, mimicking extracellular matrices such as polymers and peptide/protein hydrogels^5–9^. Also, biomaterial-based 3D cell culture platforms find several other applications including artificial organ development, organ-on-a-chip, and modeling human cancers in a test tube^10–12^. For all these applications, developing suitable biomaterials [mimicking extracellular matrix (ECM)] that potentially recapitulate the native microenvironment of tissue is most important. For example, reversible peptide/protein hydrogels, which can be easily tuned in terms of stiffness, degradability, and their ability to bind with cell adhesion sites, can facilitate the 3D culture of multiple cell types for various applications including the development of organoids. However, despite several efforts, most of the available 3D cell culture models lack proper tumor microenvironment; produce heterogeneous tumor spheroids, time-consuming protocols, and scaling these materials with exact reproducibility is difficult.

In this study, we present a class of biomaterials based on non-toxic, designed amyloid fibrils, which make thixotropic hydrogels for high-throughput *in vitro* tumor modeling. Amyloid fibrils are highly ordered protein/peptide aggregates consisting of a cross-β-sheet rich structure^13^. Amyloids are originally implicated as cytotoxic protein materials (associated with cell death in Alzheimer’s and Parkinson’s)^14^, several studies, however, indicated that many lower organisms utilize the amyloid fibrils as their ECM^15,16^. Amyloids also exhibit native functions in host organisms such as matrix formation in melanosome biogenesis^17^ and storage of hormones in secretory granules^18^. Irrespective of primary sequence, amyloid fibrils of many proteins/peptides also showed robust cell adhesion and hydrogels composed of amyloid fibrils can promote stem cell differentiation to neurons^16, 19–24^. Many advantages that distinguish amyloid hydrogels from other peptides/protein-based hydrogels, include i) cross-β structure of amyloid network possessing a unique combination of hydrophobic and hydrophilic surfaces, with an ability to bind to various small molecules and biomolecules with a high affinity^25–28^ ii) they are highly stable and resistant to extreme conditions, such as pH, temperature and proteases^29, 30^. In addition to these advantages, our designing capability rendered simple, short amyloid-forming sequences that provide amyloid biomaterials of low cost and greater ease of synthesis for several applications.

Here, we demonstrate that amyloid hydrogels designed from a highly aggregation-prone sequence of amyloid proteins and peptides provide cues for cell adhesion, proliferation, and migration. These properties help to form a cellular aggregation when various cancer cells were cultured using 3D cell culture. Further, using a simple drop cast method in 96-well plates, these hydrogels support the formation of single and uniform sized spheroid per well of various cell types possessing genotypic and phenotypic characteristics mimicking the native tumor. The spheroids formed in these hydrogels showed increased drug resistance as compared to 2D cell culture. Interestingly, these biomaterials support tumor spheroid formation from cells directly isolated from mouse xenograft breast cancer tissue, which not only overexpressed breast cancer biomarkers but also maintained the CD44^+^/CD24^-^ cell population as of isolated tumor of mouse xenograft. The present study, therefore, provides a cost-effective and simple class of biomaterials for the production of tumor spheroids to evaluate anticancer drug testing.

## Results

### Design and characterization of amyloid hydrogels for developing 3D tumor spheroids

To evaluate amyloid fibril-based hydrogels for 3D tumor spheroid formation *in vitro*, we selected a series of amino acid sequences based on the aggregation-prone region of various proteins and peptides. The highest aggregation-prone sequence was identified using TANGO for the nanofabrication of hydrogels (Fig.1a). Detailed chemical structure and sequence of selected peptide sequences are shown in Supplementary Fig.1. The designed peptides were synthesized by Fmoc-solid-phase peptide synthesis method and confirmed using ESI-MS spectroscopy (Supplementary Fig. 2). Subsequently, these peptide sequences were reconstituted for hydrogel formation. The gel formation has been confirmed by the traditional gel inversion test (Fig. 1b) and showed negligible cytotoxicity with MCF7 cells (Fig. 1c). Transmission electron microscopy (TEM) images indicated that these hydrogels are made up of nanofibrils (Supplementary Fig. 3a). The cryo-scanning electron microscopy (cryo-SEM) of these hydrogels also exhibited the presence of nanofibrils and the porosity of these hydrogels ranges from 10-18 μm, which may allow cells and nutrients to easily pass through (Fig. 1d and Supplementary Fig. 3b). The rheological data including stress-strain rheology measurement showed that all these hydrogels are soft and thixotropic (Supplementary Fig. 3c and d) as shown previously for many amyloid hydrogels^16, 21–24, 31^. The FTIR data of dried hydrogel fibrils showed cross-β-sheet rich structure (Supplementary Fig. 4a) along with ThT and CR binding suggested that the fibril networks of these hydrogels are amyloidogenic in nature (Supplementary Fig. 4b and c).

**Figure 1.**
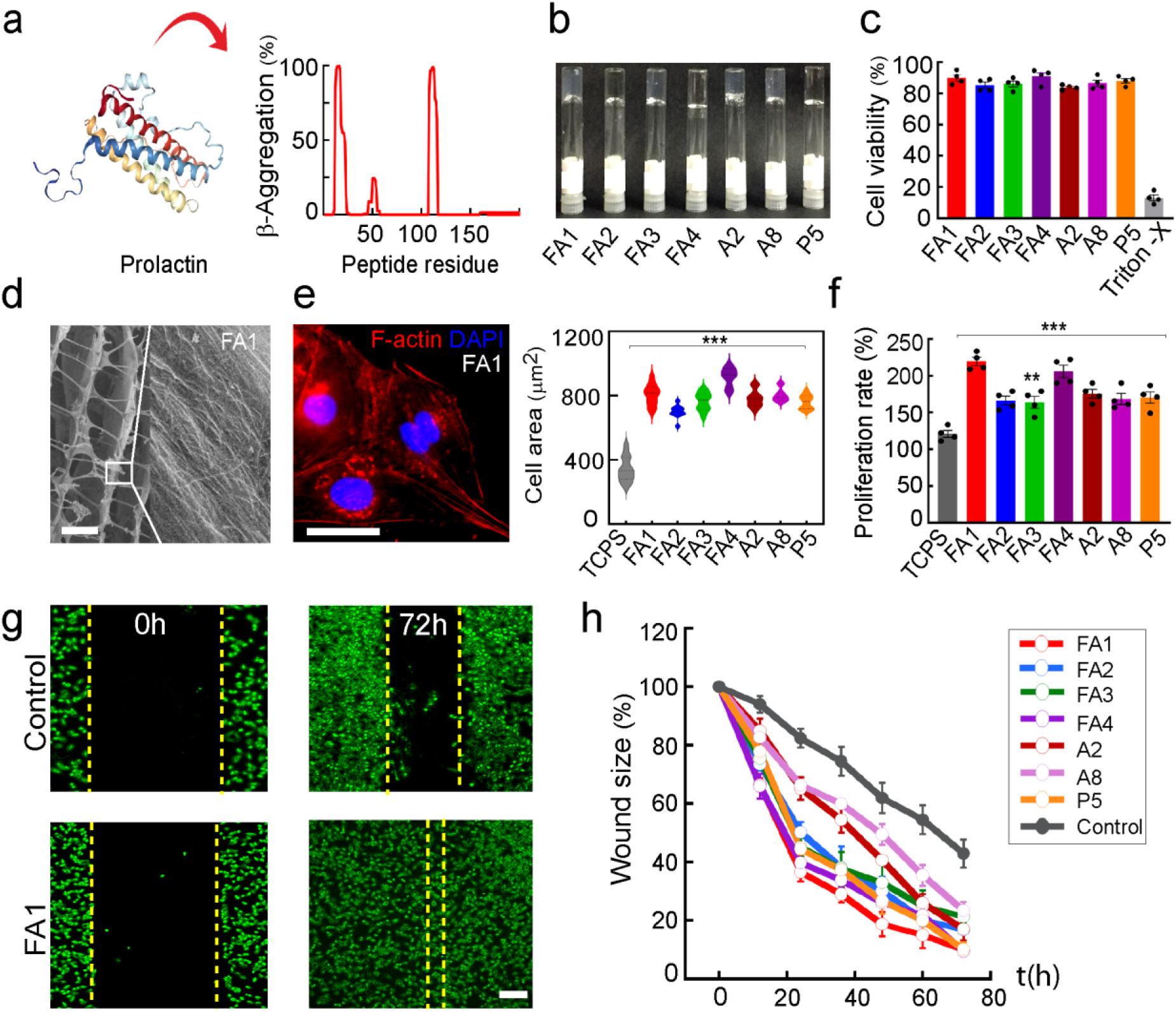
Characterization of the amyloid hydrogels as a 3D tumor scaffold system. (a) Schematic depicting designing of amyloid hydrogels from amyloidogenic proteins. TANGO plot showing the β-aggregation prone regions of prolactin as an example for selecting the amyloid peptide sequence. (b) Gel inversion test showing hydrogel formation of the designed peptides (c) Cell viability study using MTT assay of MCF7 cells cultured on the different amyloid hydrogels with >80% cell survival (n=4). Triton-X was used as a positive control. (d) Cryo-SEM images of FA1 hydrogel showing porous scaffold for hydrogel formation. The scale bar is 10 μm. (e) Immunostaining of F-actin (phalloidin, red; DAPI, blue) of MCF7 cells (left panel) cultured on hydrogel FA1 exhibit a higher cell spread area (right panel) as compared to cells cultured on tissue culture plates (TCPS). This suggests robust cell adhesion on amyloid hydrogels. The scale bar is 20 μm. (f) The proliferation of MCF7 cells by MTT assays show higher proliferation on amyloid hydrogels after 48 h compared to TCPS (n=4). (g, h) Cell migration by *in vitro* scratch assay on FA1 amyloid hydrogel (n=3). Data showing faster wound closure on FA1 hydrogel as compared to the control with time (24-72 h). The scale bar is 100 μm. Cells were pseudocoloured as green for better visualization. The values were plotted representing the mean ± s.e.m, n=4 independent experiments. Statistical significance (***p ≤ 0.001, **p ≤0.01) is determined by one-way ANOVA followed by Dunnett’s multiple comparison *post-hoc* test.

### Amyloid hydrogels support cell adhesion, proliferation, and migration of cancer cells leading to tumor spheroid formation

To support *in vitro* tumor spheroid formation, a scaffold should allow favorable cell adhesion, proliferation, and migration^8^. We tested these properties of amyloid hydrogels using the MCF7 cell line. The increased area of cell spreading and actin filament organization suggests robust cell adhesion on the amyloid hydrogels compared to the control (Fig. 1e and Supplementary Fig. 5). Further, these hydrogels also promote cell proliferation as shown by MTT assay (Fig. 1f) and cell migration as observed by wound healing assay (Fig. 1g, h and Supplementary Fig. 6) suggesting that these macroporous hydrogels allow favorable conditions for *in vitro* tumor spheroid formation.

We then performed the 3D cell culture (Fig. 2a) using various types of cancer cells [MCF7, MDA MB 231 (human breast cancer cell line), HepG2 (liver hepatocellular cells) HeLa (cervical cancer cells), and A549 (lung cancer cells)]. Along with seven different amyloid hydrogels, Matrigel was used as a control. All the five cancer cells cultured on FA1 hydrogel showed migration towards each other to form loose aggregates (day 1), which eventually transformed into larger aggregates on day 3 (Fig. 2b). From day 5 onwards, compact and spherical cellular aggregates were seen (Fig. 2b). The morphometric shape analysis data support circularity (sphericity) of the cellular aggregates was increased to ~0.9 on day 5, which then remained consistent until day 7 (Supplementary Fig. 7a). Interestingly, non-tumorigenic epithelial cells of MCF 10A and non–amyloidogenic gels Fmoc-Phe-Phe and Fmoc-Tyr-Ala gels did not show any cellular aggregation (Supplementary Fig. 7b). The spheroid diameter of five different cancer cells on FA1 amyloid hydrogel was calculated and all of them showed an increase in the spheroid size with days (Fig. 2c). Bright-field images of the day 7 spheroids by all the five cancer cells cultured on all the amyloid hydrogels suggest irrespective of different sequences, amyloid hydrogels promote cellular aggregation and 3D tumor spheroid formation by different types of cancer cells (Supplementary Fig. 7c) however, with variable size as suggested by diameter calculation (Supplementary Fig. 8). The larger spheroid sizes were in the range of 325-450 μm at day 7 and MCF7 cells formed the largest spheroids in FA1 gel (Fig. 2c). Moreover, we also observed that the size of the spheroid formed by different cells varied depending on the amyloid hydrogels used for the 3D culture. Further, the cell viability was tested for MCF7 spheroids on FA1 gel at various time points using Calcein AM/Ethidium homodimer-1 (live/dead assay) followed by fluorescence imaging. Uptake of Calcein AM resulting in green fluorescent signal indicated the viability of spheroids (Fig. 2d). Moreover, larger spheroids (>250 μm) on day 5 showed red fluorescence signal at the center (Ethidium homodimer-1 staining) suggesting a necrotic core reminiscent of the tumor^32–34^. Similar outcomes were also observed for the other four cancer cells (MDA MB 231, HeLa, HepG2, and A549) cultured on FA1 gel (Supplementary Fig. 9a). This is further supported by the 3D surface plot of MCF7 spheroids on day 7 showing a clear distribution of live and dead cells corresponding to three major zones of proliferation, quiescent and necrotic zones (Fig. 2d, Supplementary Fig. 9b). The larger spheroids containing a dead core at the center could be due to a decrease in gaseous exchange resembling cellular heterogeneity of the solid tumor^32, 33, 35^.

**Figure 2.**
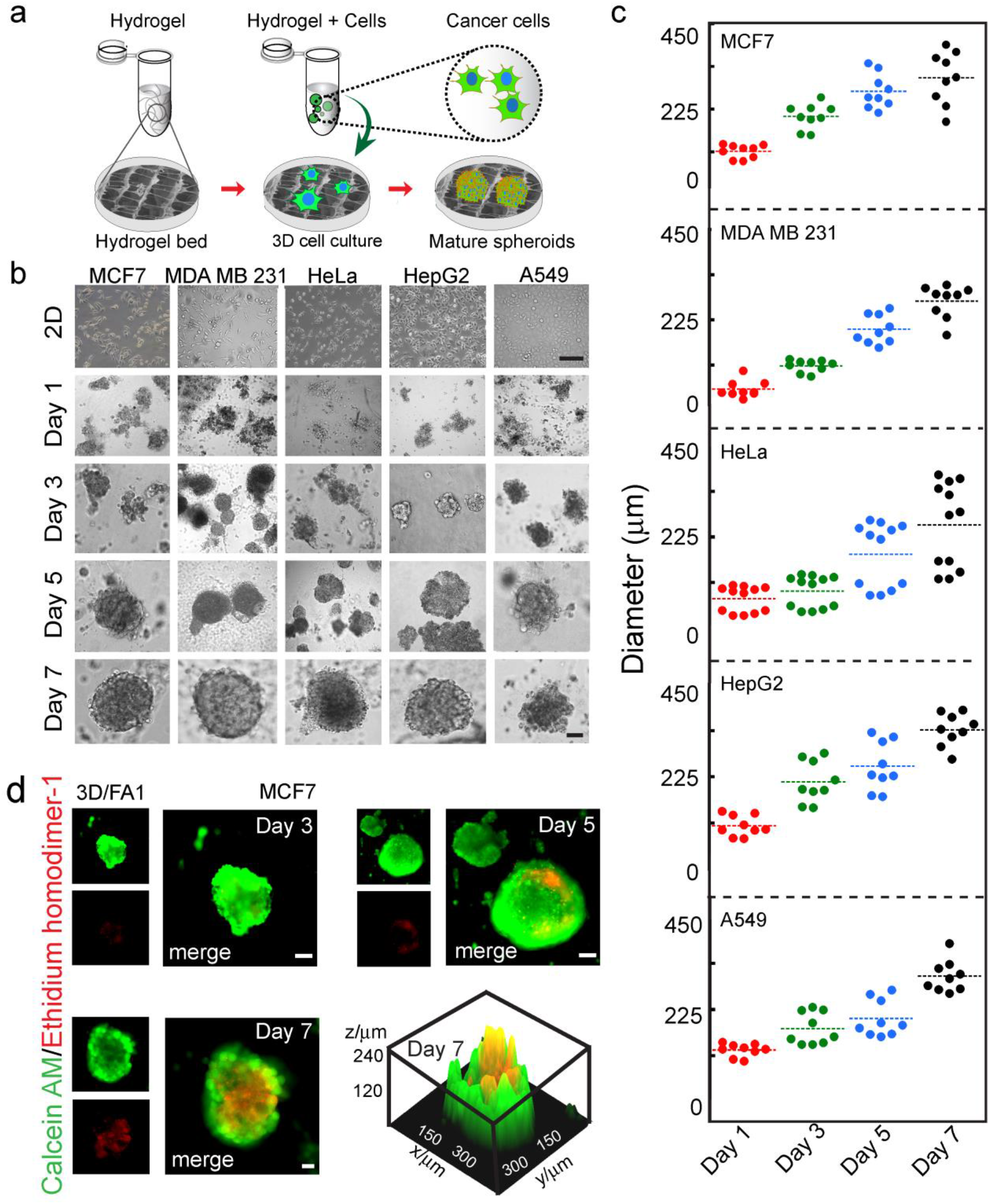
Three-dimensional (3D) cell culture showing cellular aggregation by different cancer cells on FA1 amyloid hydrogel. (a) Schematic showing 3D cell culture of cancer cells with amyloid hydrogel. A mixture of gel and cells was spread on gel-coated culture plates and were incubated for 7 days. (b) Bright-field images showing cellular aggregation over time by five different cell types on FA1 amyloid hydrogel. The scale bar is 200 μm. The data are showing a gradual formation of heterogeneous size spheroids with time. (c) Corresponding growth kinetics was quantified, which showed an increase in the diameter of cell aggregates over time. (d) Cell viability assay of spheroids using Calcein AM/Ethidium homodimer-1 staining showing the viable cells at the early stage followed by the appearance of the necrotic core inside the large spheroid at the later stage. The 3-D surface plot at the day 7 spheroid showing the necrotic core at the center of cellular spheroids.

### Expression of cancer biomarkers support spheroid formation with amyloid hydrogels

We further evaluated the cancer biomarker genes by RT-qPCR in these spheroids formed using 3D cell culture and Matrigel was used as control. Genes selected for the analysis were associated with epithelial to mesenchymal transition (EMT), cell proliferation, migration, cancer stem cells, cell cycle, and adhesion (Supplementary Table 1), which are known to be overexpressed in cancers, such as breast, lung, and ovarian^36, 37^. We observed upregulation of the pro-oncogenic markers such as ERBB2, CDH1, SLUG, CTNBB1, VEGF, CD44, CDC 20, and ITGB4 genes. In contrast, tumor suppressor genes CCND2 and CD24 were downregulated as compared to 2D monolayer culture (Fig. 3a, Supplementary Fig. 10). Further, an immunofluorescence study with day 5 spheroid suggested higher E-cadherin expression in MCF7, HepG2, and A549 spheroids formed in FA1 gel as compared to 2D cell culture [Fig. 3b (upper panel), c and Supplementary Fig. 11a]. The higher E-cadherin expression is further confirmed using Western blot of MCF7 spheroids (Fig. 3d).

**Figure 3.**
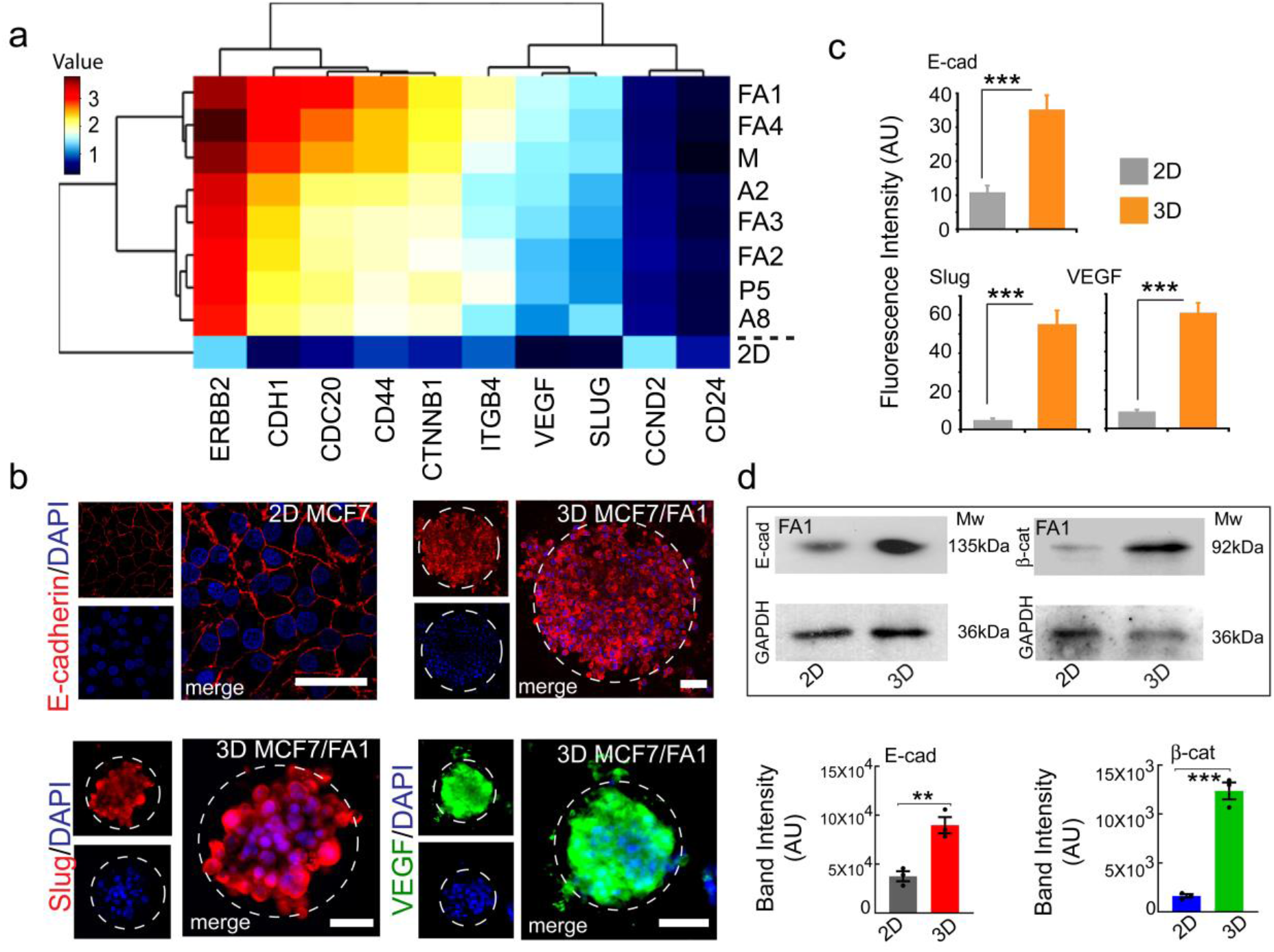
Expression of the cancer biomarkers in MCF7 spheroids cultured with amyloid hydrogels. (a) Heat map illustrating hierarchical clustering of RT-qPCR data of cancer gene expression in 3D MCF7 spheroids compared to Matrigel (positive control) and 2D cell culture. Red indicates higher and blue indicates lower expression of the genes compared to the 2D monolayer cells. (b, c) Immunofluorescence microscopy showing higher expression E-cadherin (day 5 spheroid), EMT markers of Slug, and VEGF (day 7 spheroid) of MCF7 cells compared to corresponding 2D culture. DAPI was used for staining the nucleus. (d) Western blot showing E-cadherin (day 5) and β-catenin (day 7) over-expression in the 3D spheroid of MCF7 cell compared to 2D monolayer culture on FA1 amyloid hydrogel. Statistical significance (**p ≤ .01, ***p ≤ .001) is determined by one-way ANOVA followed by Dunnett’s multiple comparisons *post-hoc* test.

Similarly, higher N-cadherin and β1-integrin expression were observed for HeLa, and MDA MB 231 cells (as these cells are E-cadherin negative^38^) (Supplementary Fig. 11a). To directly probe the role of cell adhesive protein for cellular aggregation, the 3D culture was done with EGTA supplemented media (to remove Ca^2+^, important for adhesion protein to function^39^). No cellular aggregation and spheroids formation was observed, suggesting an essential role of calcium-dependent adhesion protein such as cadherin for spheroid formation *in vitro* (Supplementary Fig. 11b).

Interestingly, the spheroids also over-express EMT markers (Slug and VEGF) and β-catenin on prolonging incubation (day 7) as shown using immunofluorescence study and Western blotting, respectively (Fig. 3b, lower panel, 3c, d). The data suggest that after long incubation, these spheroid undergo EMT-like transitions similar to the tumor *in vivo*^40^.

### High-throughput single spheroid formation in 96-well formats using drop cast method

To make a reproducible single spheroid in 96-well format, we developed a new methodology, where hydrogel and cell mixture was cast as a drop on top of the corresponding preformed hydrogel bed in 96-well plate. We termed this as the drop-cast (DC) method (Fig. 4a). To examine the formation of a single spheroid per well, five cancer cell lines and seven amyloid hydrogels were cultured using the drop cast method for 5 days (Fig. 4a, Supplementary Fig. 12a). We observed an increase in drop size along with a darker appearance of the spheroid with uniformity in shape and smooth/defined boundaries for all cells in each of the gels under study (Fig. 4b). The darker appearance is due to the compact cell aggregation with time^41, 42^. Since the drop cast method maintains the spherical shape throughout the incubation period (Supplementary Fig. 12b, left panel), cell density (solidity) measurement with time was done^42^. The data showed the solidity increased from 0.7 (day 1) to 0.95 (day 3) and then remained stable till day 5 (Supplementary Fig. 12b, right panel). Further, the cell viability and the necrotic core formation of MCF7 spheroids formed by this method in various amyloid hydrogels (day 3) was examined using Calcein AM/Ethidium homodimer-1 staining. MCF7 spheroids formed in all amyloid hydrogels showed cell death at the core suggesting the formation of the necrotic zone, however, with varied size (1 mm to 3.4 mm) in various hydrogels. The necrotic core size of MCF7 spheroids in FA1 and FA4 hydrogel were comparatively less as compared to other gels and Matrigel (Fig. 4c, d).

**Figure 4:**
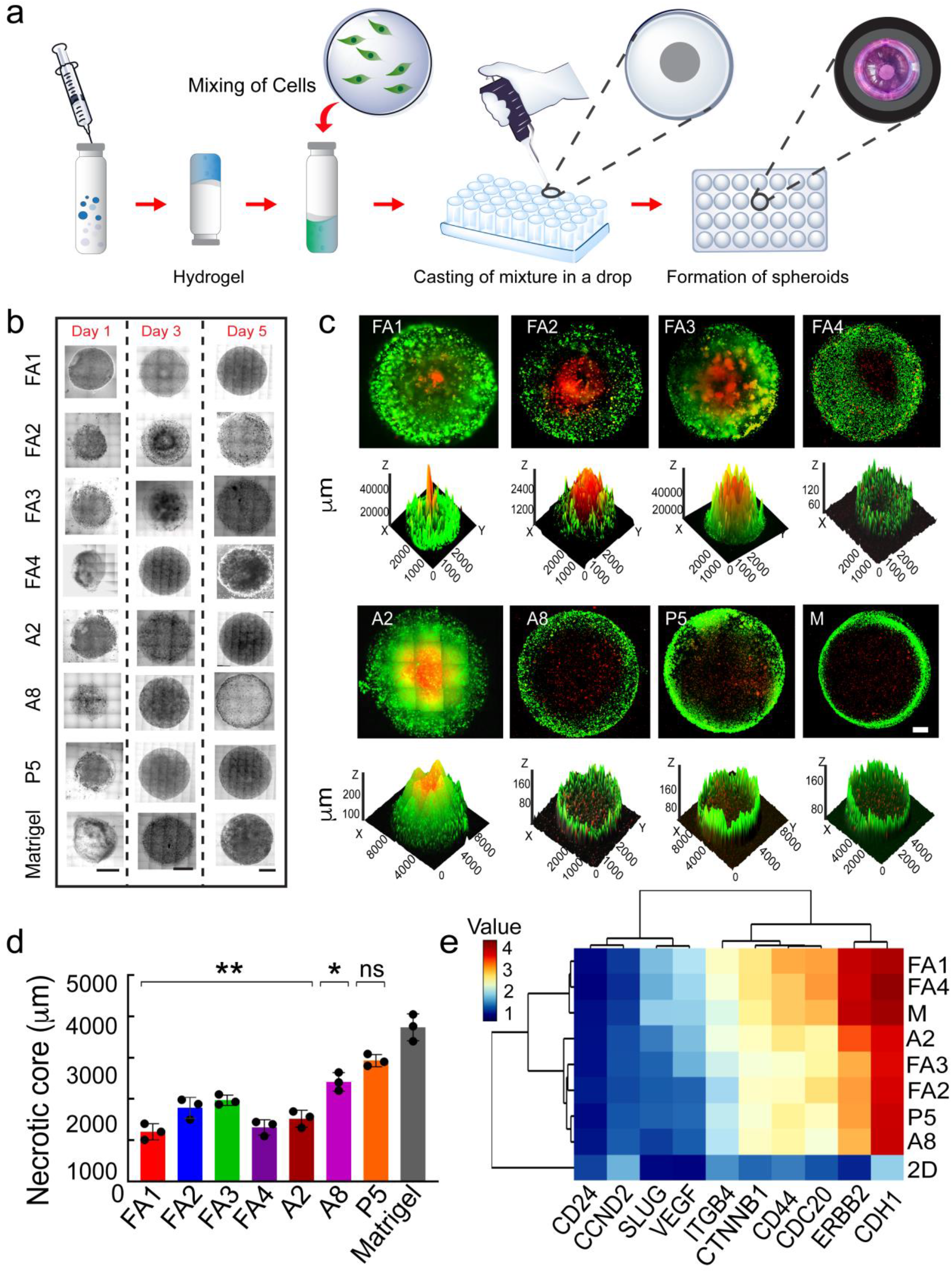
High-throughput single spheroid formation by MCF7 cells using drop cast method. (a) Schematic showing steps for single spheroid generation of MCF7 cells cultured with various amyloid hydrogels using the drop cast method. (b) Bright-field images showing spheroid growth by MCF7 cells with time using various amyloid hydrogels. Increment of both the spheroid size and darker appearance indicating denser cellular packing over time. Scale bars are 500 μm. (c) Confocal imaging with Calcein AM (green, live cells)/Ethidium homodimer-1 (red, dead cells) staining with day 3 spheroid of MCF7 cells showing necrotic core. Corresponding 3D construction plot indicating that dead cells were located in the central region of the spheroid. The scale bar is 500 μm. (d) Quantification of necrotic core size of MCF7 spheroids in different amyloid hydrogels showing the smaller necrotic region of FA1 and FA4. (e) Heat map illustrating the hierarchical clustering of RT-qPCR data of gene expression in MCF7 3D spheroids in various hydrogels compared to corresponding 2D cell culture. The fold changes were presented as log2. M denotes Matrigel. Data represent three biological repeats and are displayed as mean ± s.e.m and statistical significance (*p ≤ .05, **p ≤ .01, ***p ≤ .001) is determined by one-way ANOVA followed by Dunnett’s multiple comparison test.

### Upregulation of oncogenic markers in drop cast tumor spheroids analyzed by RT-qPCR and microarray

To analyze the expression levels of cancer gene markers in spheroid using the drop cast method, RT-qPCR analysis was done for the MCF7 spheroids (day 3). Similar to 3D cell culture, the drop cast spheroids also showed upregulation of pro-oncogenic markers such as CDH1, CTNBB1, SLUG, CD44, ERBB2, ITGB4, CDC20, and VEGF and downregulation of CCND2 and CD24 as compared to 2D monolayer culture (Fig. 4e, Supplementary Fig. 13). The heat map further showed that the gene expression pattern of amyloid hydrogel spheroids (in FA1 and FA4) was similar to the Matrigel spheroids (Fig. 4e).

Further, genome-wide gene expression analysis was performed by microarray using MCF7 spheroids (day 3) cultured with FA1, FA4, A2 along with Matrigel as control. The total number of altered genes in FA1, FA4, A2, and Matrigel were 5240, 5426, 7558, 5191, respectively (Supplementary Table 2). Further, to understand the presence of breast cancer marker genes involved during spheroids formation, signature biomarker genes were sorted from the complete human cancer array^43^. Heat map for the presence of signature marker breast cancer genes in the spheroids indicated upregulation of pro-oncogenes and downregulation of the tumor suppressor genes including pro-apoptotic genes (Fig. 5a and Supplementary Fig. 14a). This suggests 3D MCF7 spheroids can mimic the tumor microenvironment by modifying the key cellular pathways involved in tumor formation. The significantly affected gene functions were sorted in terms of biological processes, cellular components, and molecular functions, we observed that affected genes were associated with the inflammatory response, apoptosis, cell migration, growth, proliferation, and adhesion using Metascape analysis (Fig. 5b, left panel and Supplementary Fig. 14b) as expected in tumor formation^44^. For example, the higher expression of MYC, KRAS, EGFR, VEGF, and WNT genes, are known to be responsible for tumor cell growth, proliferation, metabolism, and apoptosis in breast cancer^45^ (Fig. 5b, right panel).

**Figure 5.**
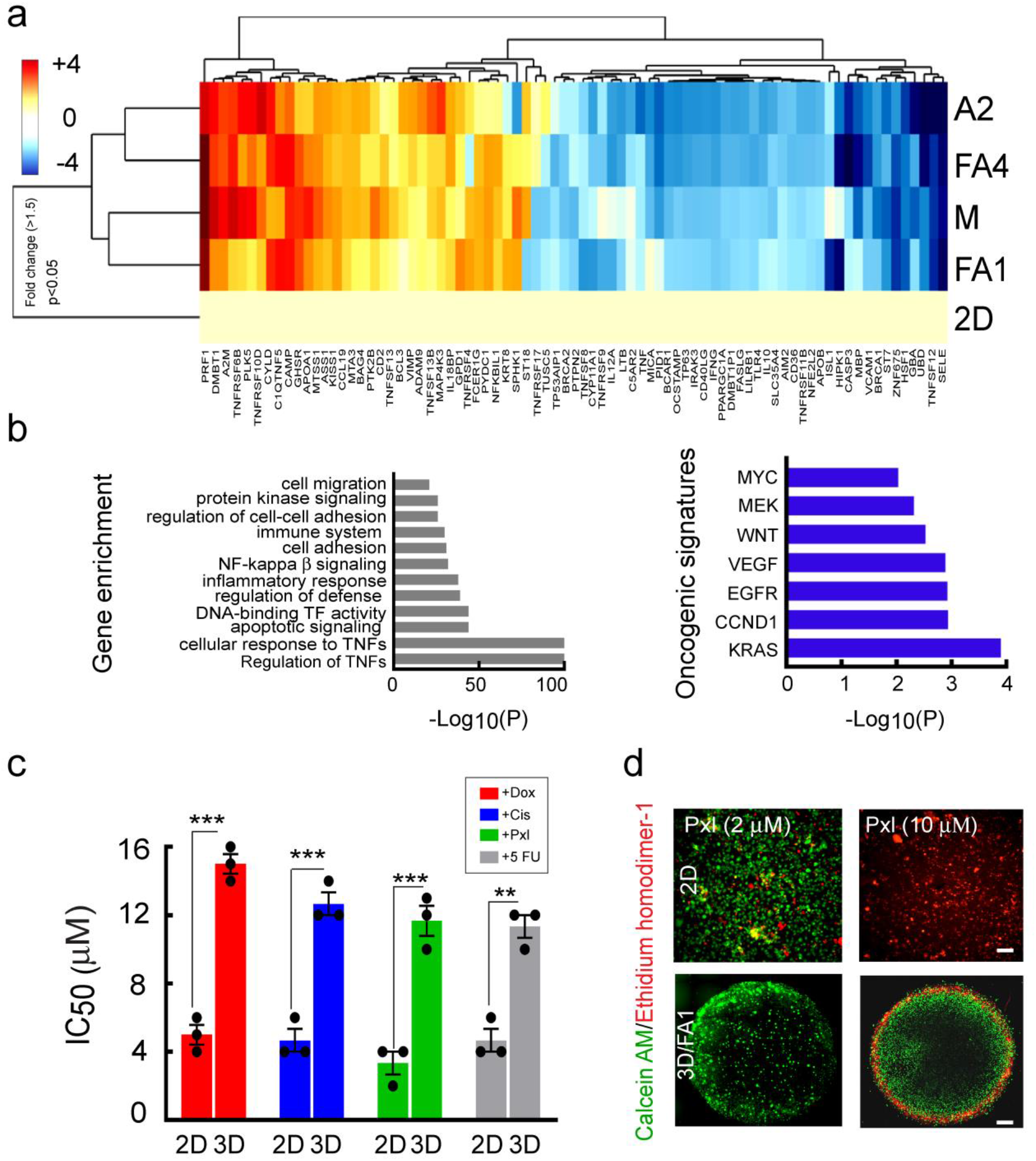
Genome-wide cancer gene expression study of 3D MCF7 spheroids cultured with amyloid hydrogels using microarray. (a) Heat map illustrating a comparative study of breast cancer biomarker genes in 3D MCF7 spheroids (drop cast method). Matrigel (M) is used as a control. The fold changes are presented as log2 to visualize the heat map. (b) Gene ontology enrichment analysis for the differentially regulated genes in 3D spheroid showing various biological processes altered during spheroid formation. Metascape analysis for oncogenic signatures in 3D spheroid showing over-expression of breast cancer genes like MYC, KRAS, EGFR, and VEGF (right panel). (c) Toxicity assay using Annexin V/PI staining followed by FACS analysis showing various cancer drug resistance (higher IC_50_ value) by 3D MCF7 spheroid in comparison to 2D culture. (Dox is Doxorubicin; Cis is Cisplatin, Pxl is Paclitaxel and 5FU is 5-Fluorouracil). The values plotted represent mean ± s.e.m, n=3 independent experiments. ***p ≤ 0.001, **p ≤ 0.01) (d) The spheroid staining with Calcein AM (green, viable cells) and Ethidium homodimer-1 (red, dead cells) post drug treatment showing that the cell death is restricted to outer cell layers compared to complete cell death in 2D cell culture. This indicates that the compact structure of the 3D spheroid does not allow the drug to penetrate.

Subsequently, gene enrichment analysis indicated that the statistically significant regulatory pathways such as NFκB1, STAT3, JUN, TP53, RELA, and SP1 were responsible for the induction of tumor spheroid formation (Supplementary Fig. 14c). Further, the NFκB1 regulatory pathway showed the highest log(P) value, suggested that this might be the central pathway for the induction of the amyloid hydrogel spheroids. These findings agree with previous studies on NFκB1 regulatory pathway in breast cancer^46^. The data, therefore, imply that tumor spheroid formation using amyloid hydrogel mimics the tumor formation *in vivo* in terms of gene expression.

### Drug resistance of 3D cancer cell spheroids

We further evaluated the efficacy of various chemotherapeutic drugs (cisplatin, doxorubicin, paclitaxel, pentostatin, and 5-fluorouracil) against cells cultured in 2D versus drop cast spheroids (day 3) by MTT assay (Supplementary Fig. 15). Drop cast spheroids showed much higher resistance to all five chemotherapeutic drugs as compared to the cells in 2D culture (two-fold high IC_50_ for spheroid versus 2D cells). This is further supported by the quantitative cell death assay using Annexin V/PI followed by FACS analysis (Fig. 5c, Supplementary Fig. 16). These findings confirm that drug penetration is hindered in the dense spheroids in 3D culture, which demonstrated the lesser sensitivity of the drugs for the cells in 3D spheroids as compared to 2D monolayer culture. Indeed, the Calcein AM/Ethidium homodimer-1 staining suggested that cell death in 3D spheroid are less and restricted to outer cell layers as compared to extensive cell death in 2D culture (Fig. 5d and Supplementary Fig. 17).

### Tumor spheroid from breast tumor tissues of xenograft mice

To directly demonstrate that tumor spheroids can be developed from *in vivo* cancer tissue with heterogeneous cells, we developed tumor xenograft in SCID mice by injecting MCF7 cells (Fig. 6a). After characterizing tumor tissue by hematoxylin and eosin (H&E) staining (Fig. 6b), tumor tissues were processed into single-cell suspension^47^, which has a heterogeneous cell population (Fig. 6c). Similar to various cancer cell lines (Fig. 2b and 4b), these tumor-derived cells also showed time-dependent cellular aggregation in 3D cell culture in FA1 and FA4 amyloid hydrogels along with matrigel as a positive control (Fig. 6d, Supplementary Fig. 18a). When these cells were drop-casted using FA1 and FA4 hydrogels, spheroids were formed within 3 days with a proliferative outer region (Fig. 6d, Supplementary Fig. 18a). Staining by Calcein AM/Ethidium homodimer-1 of the tumor spheroids indicated that cells were viable and formed a necrotic core (Fig. 6d and 6e). The gene expression pattern of cancer markers in these tumor spheroids was similar to the original tumor suggesting that the drop cast tumor spheroid recapitulates the features of the original tumor (Fig. 6f). This is further supported by a similar cell population of CD44^+^/CD24^-^ cells (cancer stem cell markers population^48^) at day 5 spheroids (Fig. 6g and Supplementary Fig. 18b). Additionally, these tumor spheroids also showed high resistance against doxorubicin as IC_50_ for 3D spheroid was significantly higher compared to the 2D (Fig. 6h, Supplementary Fig. 18c).

**Figure 6.**
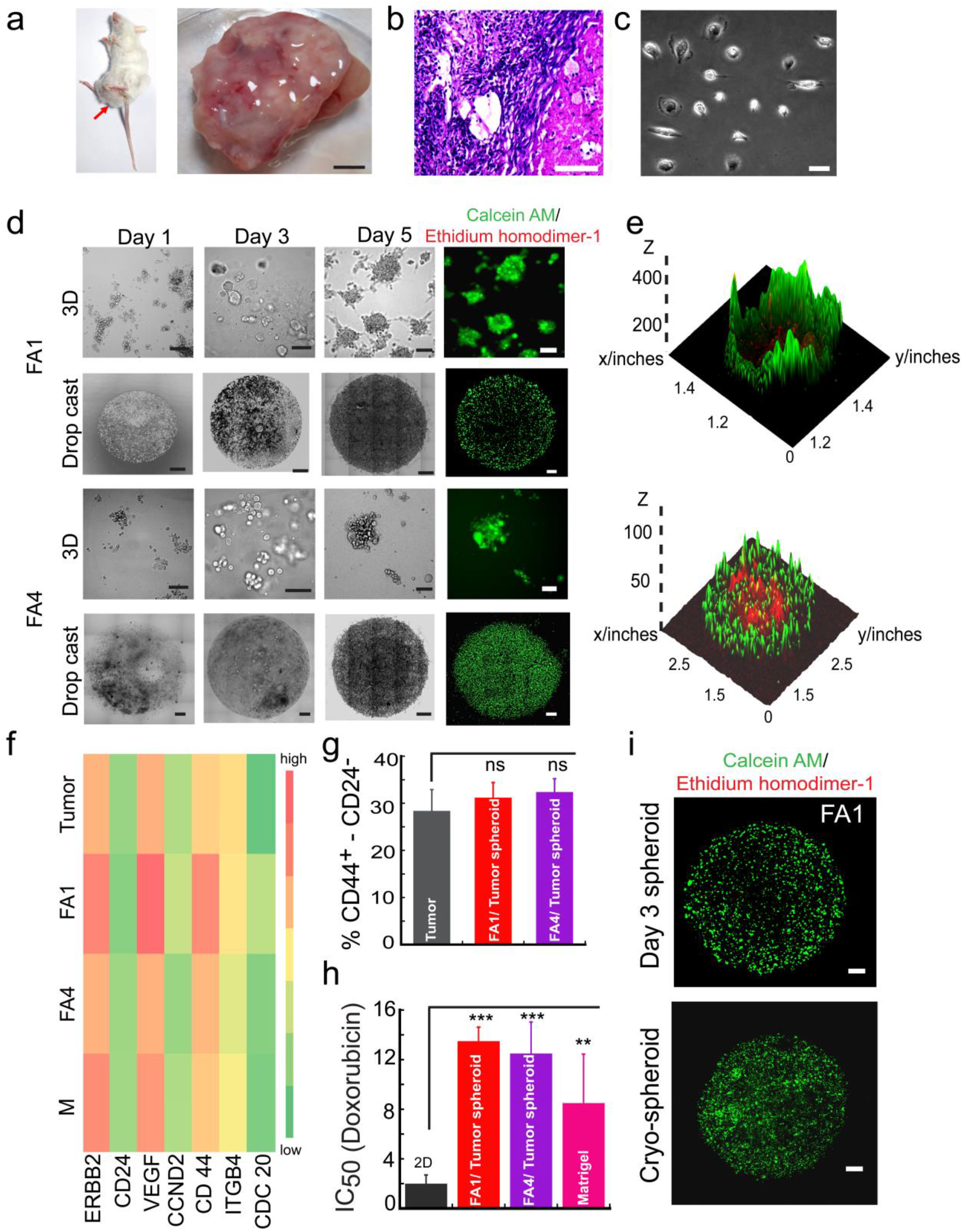
Amyloid hydrogels support tumor spheroid formation of cells derived from tumor tissue of xenograft mice. (a) Xenograft mouse with MCF7 cell injection showing tumor formation after a week (left panel). Representative image of isolated tumor after the sacrifice of an animal (right panel). The scale bar is 10 mm. (b) H&E staining of tumor tissue showing the hyperproliferative nature. The scale bar is 50 μm. (c) Morphology of the cells isolated from tumor tissue for the development of 3D spheroid. The scale bar is 100 μm. (d) Formation of tumor spheroids using 3D cell culture and drop cast method with FA1 and FA4 hydrogel over time. The scale bar for the 3D cell culture is 100 μm and for the drop cast method is 500 μm. (e) Calcein AM (green, live cells), Ethidium homodimer-1 (red, dead cells) staining followed by microscopy imaging showing the viability of the cells in the spheroids. The corresponding 3D surface plot of tumor spheroids with FA1 and FA4 at day 5 by Image J showing proliferating viable rim and necrotic core formation at the center. (f) Heat map showing similar expression levels of cancer genes in 3D drop cast spheroid and original tumor isolated from the mouse. (g) Percentage of CD44^+^/CD24^-^ cell population estimated by FACS sorting showing similar population CD44^+^/CD24^-^ cells in drop cast spheroid (day 5) and the original tumor sample. (h) Efficacy of anti-cancer drug doxorubicin on 3D breast tumor spheroids showing significant resistance (3 fold higher IC_50_) as compared to 2D cultured cells. Three biological repeats were done and mean ± s.e.m. values were plotted. Statistical significance (*p ≤ .05, **p ≤ .01, ***p ≤ .001, ****p ≤ .0001) is determined by one-way ANOVA followed by Dunnett’s multiple comparison test. (i) Calcein AM/Ethidium homodimer-1 staining of breast tumor spheroids showing the viability of cells in spheroids before and after two months of cryopreservation. The scale bar is 500 μm.

Next, we asked whether these tumor spheroids can be cryopreserved for drug testing and genetic analysis. Cryo-preserved spheroid showed high cell viability (Fig. 6i) suggesting ready-to-use tumor spheroid can be developed with amyloid hydrogel for biobanking of cancer tissues.

## Discussion

Cancer prognosis is a major challenge particularly due to the heterogeneity, architecture, and genotype-phenotype variations among the patients for different cancer types^49,50^. Thus, the failure rate of anticancer drugs during clinical trials is higher than any other therapeutics. This is mainly due to the absence of reliable preclinical models for the assessment of the drugs (particularly for personalized medicine). In this essence, biomaterial based 3D cell culture systems or *in vitro* organoid models have gained interest for clinical outcome and hence are becoming more prominent in drug discovery^9, 51^.

Numerous systems have been developed for making tumor spheroid or organoid *in vitro* using polymeric biomaterials^52^, protein/peptide hydrogels^53^, and also using various methods of producing spheroids such as hanging drop^1, 54^, and spinning flask^55^. But most of these are either lack of achieving homogeneous tumor spheroid formation consistently and/or unable to mimic microenvironments of original tumor^56, 57^. A reliable, inexpensive, and tunable synthetic matrix mimicking the extracellular matrix of tissue for tumor spheroid development is still lacking.

In this study, we describe self-assembled amyloid fibril-based hydrogels, which provide the necessary structural and biochemical support for the formation of tumor spheroids from various cancer cell lines as well as cells from tumors. Amyloid fibrils are well known for their cell-adhesive nature^16,20,58^ and were previously used for developing hydrogel for stem cell differentiation^20,24,59^. These amyloid hydrogels are generally made from short peptide sequences, which are easy to synthesize, scalable, and cost-effective. The soft nature of the hydrogel along with thixotropicity allowed an easy cell modeling with gel solution for 3D cell culture for tumor spheroid and/or possible organoid technology^31, 60^. Further, the unique surface of the amyloid hydrogel provides a favorable microenvironment for cell adhesion, survival, and proliferation, while also providing mechanical support to the cells for 3D tumor spheroid formation (Fig. 1).

However, similar to other polymeric/peptide hydrogels^6,53^, these amyloid hydrogels also promote cellular aggregation and spheroid formation but with heterogeneity in size (Fig. 2). Moreover, depending on the peptide sequence of the amyloid hydrogels, the variability of spheroids size was also observed as different peptide sequences and porosity of the hydrogel network might influence cell adhesion and migration of cells^61^. Further, irrespective of cell and gel types, once the spheroid is large enough, it showed a necrotic core reminiscent of tumor *in vivo*^32–34^. All these 3D cultured spheroids also showed gene expression similar to cancer tissue as well as exhibited a higher drug resistance as compared to 2D culture (Fig. 3, and 5). This observation raises the exciting possibility for 3D tumor spheroid modeling *in vitro* for possible cancer therapeutic application. However, important to note that the use of 3D culture for high-throughput screening assays is limited due to the difficulty in producing homogeneous and reproducible size spheroid cultures^62^. Although the variability in spheroid size can be overcome by various methods such as hanging drop^54^. Therefore, achieving a close mimic of the tumor microenvironment often gets difficult due to the lack of biomaterials in this tumor spheroid development. To overcome all these limitations, we further developed a “drop-cast” method capable of generating homogenous spheroids for high-throughput drug assays (Fig. 4). This procedure consistently generated single spheroids with reproducible size in each well of 96-well plate within 3 days of culture time with proliferating cells in the rim. These spheroids also maintained the tumor microenvironment by upregulating pro-oncogenes and downregulating the tumor suppressor genes (Fig. 5) and showed higher cancer drug resistance compared to 2D culture. However, the cellular environment in tumors significantly differs from the cancer cell lines as multiple cellular systems (cellular heterogeneity) exists in tumors *in vivo*^50^. The presented hydrogels also support tumor spheroid formation with mouse xenograft cancer tissue, which not only maintained the cancer gene expression but also cancer stem cell population, similar to the original tumor (Fig. 6).

Further, restricted interstitial transport is a major *in vivo* barrier for limited drug efficacy in solid tumor tissue^63^. In this aspect, it has been observed that a 3D spheroid culture model that mimics the microenvironment and diffusion processes of solid tumors achieves more accurate predictions of *in vivo* drug evaluation^34^. Indeed, the efficacy of anti-cancer drugs using the present tumor tissue-derived spheroid showed high resistance of drugs compared to 2D culture supporting the fact that drug doses estimated from these spheroids might useful for treating the tumors in patients.

To overcome the problem of preparing a 3D tumor model immediately for drug testing, cryopreservation of spheroids is an essential requirement^64^. Cryo-preserved spheroids will not only be used for personalized medicine but also during the development or repurposing of the drugs. The amyloid hydrogel supported spheroid showed high cell viability even after two months of cryopreservation further suggesting that the designed amyloid hydrogel are suitable biomaterials candidates for tumor spheroid technology (Fig. 6).

In conclusion, our data suggest that the amyloid hydrogels are one of the superior classes of biomaterials for producing tumor spheroid for cancer drug development, repurposing of drug and recapitulating the tumor property in 96-well plate for future cancer biology study.

## Acknowledgments

We acknowledge IIT Bombay central facilities for FACS, TEM, Cryo-SEM, Rheology, FT-IR, Spinning disc microscopy, and confocal microscopy for performing the experiments. We also acknowledge Genotypic Technology Private Limited Bangalore for microarray processing and verifying the data analysis reported in this publication. The authors thank the DBT-BIRAC PACE scheme for financial support. N.S. acknowledges IIT institutional fund for the fellowship. We are thankful to Prof. Shamik Sen of IIT Bombay for the kind gift of FITC-labeled CD44 and PE-CD24 antibodies.

## Author Contribution

N.S. and S.K.M. conceived the project and planned the experiments. N.S. carried out all the experiments unless mentioned otherwise. K.P. carried out the cryo-SEM, TEM, and rheology experiments, helped in analysis and reviewing the manuscript. A.N. acquired FACS data, developed mice xenograft, and also helped in reviewing the manuscript. P.K. synthesized the peptide and analyzed the corresponding mass. A.S. helped in hydrogel preparation. D.D. helped during capturing confocal images. D.C. carried out an FT-IR experiment.

## Conflict of interest

The authors declare no conflict of interest.

## Notes

### Competing Interest Statement

The authors have declared no competing interest.

## References

1. Pampaloni, F., Reynaud, E.G. & Stelzer, E.H. The third dimension bridges the gap between cell culture and live tissue. Nat. Rev. Mol. Cell Biol. 8, 839–845 (2007).

2. Hutchinson, L. & Kirk, R. High drug attrition rates—where are we going wrong? Nat. Rev. Clin. Oncol. 8, 189–190 (2011).

3. Duval, K. et al. Modeling Physiological Events in 2D vs. 3D Cell Culture. Physiology (Bethesda) 32, 266–277 (2017).

4. Day, C.-P., Merlino, G. & Van Dyke, T. Preclinical mouse cancer models: a maze of opportunities and challenges. Cell 163, 39–53 (2015).

5. Lee, K.Y. & Mooney, D.J. Hydrogels for Tissue Engineering. Chem. Rev. 101, 1869–1880 (2001).

6. Zhang, S. Fabrication of novel biomaterials through molecular self-assembly. Nat. Biotechnol. 21, 1171–1178 (2003).

7. Cushing, M.C. & Anseth, K.S. Materials science. Hydrogel cell cultures. Science 316, 1133–1134 (2007).

8. Tibbitt, M.W. & Anseth, K.S. Hydrogels as extracellular matrix mimics for 3D cell culture. Biotechnol. Bioeng. 103, 655–663 (2009).

9. Li, Y. & Kumacheva, E. Hydrogel microenvironments for cancer spheroid growth and drug screening. Sci. Adv. 4, eaas8998 (2018).

10. Lancaster, M.A. et al. Cerebral organoids model human brain development and microcephaly. Nature 501, 373–379 (2013).

11. Ingber, D.E. Reverse Engineering Human Pathophysiology with Organs-on-Chips. Cell 164, 1105–1109 (2016).

12. Clevers, H. Modeling Development and Disease with Organoids. Cell 165, 1586–1597 (2016).

13. Greenwald, J. & Riek, R. Biology of amyloid: structure, function, and regulation. Structure 18, 1244–1260 (2010).

14. Chiti, F. & Dobson, C.M. Protein misfolding, functional amyloid, and human disease. Annu. Rev. Biochem. 75, 333–366 (2006).

15. Otzen, D. & Nielsen, P.H. We find them here, we find them there: Functional bacterial amyloid. Cell. Mol. Life Sci. 65, 910–927 (2008).

16. Jacob, R.S. et al. Amyloids Are Novel Cell-Adhesive Matrices. Adv. Exp. Med. Biol. 1112, 79–97 (2018).

17. Fowler, D.M. et al. Functional amyloid formation within mammalian tissue. PLoS Biol. 4, e6 (2006).

18. Maji, S.K. et al. Functional amyloids as natural storage of peptide hormones in pituitary secretory granules. Science 325, 328–332 (2009).

19. Jacob, R.S., Das, S., Ghosh, D. & Maji, S.K. Influence of retinoic acid on mesenchymal stem cell differentiation in amyloid hydrogels. Data in Brief 5, 954–958 (2015).

20. Jacob, R.S. et al. Cell Adhesion on Amyloid Fibrils Lacking Integrin Recognition Motif. J. Biol. Chem. 291, 5278–5298 (2016).

21. Jacob, R.S. et al. Self healing hydrogels composed of amyloid nano fibrils for cell culture and stem cell differentiation. Biomaterials 54, 97–105 (2015).

22. Das, S., Jacob, R.S., Patel, K., Singh, N. & Maji, S.K. Amyloid Fibrils: Versatile Biomaterials for Cell Adhesion and Tissue Engineering Applications. Biomacromolecules 19, 1826–1839 (2018).

23. Das, S., Kumar, R., Jha, N.N. & Maji, S.K. Controlled Exposure of Bioactive Growth Factor in 3D Amyloid Hydrogel for Stem Cells Differentiation. Adv. Healthc. Mater. 6 (2017).

24. Das, S. et al. Implantable amyloid hydrogels for promoting stem cell differentiation to neurons. NPG Asia Mater. 8, e304–e304 (2016).

25. Nelson, R. & Eisenberg, D. Recent atomic models of amyloid fibril structure. Curr. Opin. Struct. Biol. 16, 260–265 (2006).

26. Maji, S.K., Wang, L., Greenwald, J. & Riek, R. Structure-activity relationship of amyloid fibrils. FEBS Lett. 583, 2610–2617 (2009).

27. Nilsson, K.P. Small organic probes as amyloid specific ligands--past and recent molecular scaffolds. FEBS Lett. 583, 2593–2599 (2009).

28. Ghosh, D. et al. Complexation of amyloid fibrils with charged conjugated polymers. Langmuir 30, 3775–3786 (2014).

29. Meersman, F. & Dobson, C.M. Probing the pressure-temperature stability of amyloid fibrils provides new insights into their molecular properties. Biochim. Biophys. Acta 1764, 452–460 (2006).

30. Mesquida, P., Riener, C.K., MacPhee, C.E. & McKendry, R.A. Morphology and mechanical stability of amyloid-like peptide fibrils. J. Mater. Sci.: Mater. Med. 18, 1325–1331 (2007).

31. Zaman, M.H. et al. Migration of tumor cells in 3D matrices is governed by matrix stiffness along with cell-matrix adhesion and proteolysis. Proc. Natl. Acad. Sci. U S A 103, 10889–10894 (2006).

32. Vinci, M. et al. Advances in establishment and analysis of three-dimensional tumor spheroid-based functional assays for target validation and drug evaluation. BMC Biol. 10, 29 (2012).

33. De Sousa, E.M.F., Vermeulen, L., Fessler, E. & Medema, J.P. Cancer heterogeneity--a multifaceted view. EMBO Rep. 14, 686–695 (2013).

34. Edmondson, R., Broglie, J.J., Adcock, A.F. & Yang, L. Three-dimensional cell culture systems and their applications in drug discovery and cell-based biosensors. Assay Drug Dev. technol. 12, 207–218 (2014).

35. Däster, S. et al. Induction of hypoxia and necrosis in multicellular tumor spheroids is associated with resistance to chemotherapy treatment. Oncotarget 8, 1725–1736 (2017).

36. Slamon, D.J. et al. Studies of the HER-2/neu proto-oncogene in human breast and ovarian cancer. Science 244, 707–712 (1989).

37. Chen, C., Zhao, S., Karnad, A. & Freeman, J.W. The biology and role of CD44 in cancer progression: therapeutic implications. J. Hematol. Oncol. 11, 64–64 (2018).

38. Velez, D.O. et al. 3D collagen architecture induces a conserved migratory and transcriptional response linked to vasculogenic mimicry. Nat. Commun. 8, 1651 (2017).

39. Kim, S.A., Tai, C.-Y., Mok, L.-P., Mosser, E.A. & Schuman, E.M. Calcium-dependent dynamics of cadherin interactions at cell-cell junctions. Proc. Natl. Acad. Sci. U S A 108, 9857–9862 (2011).

40. Valastyan, S. & Weinberg, R.A. Tumor metastasis: molecular insights and evolving paradigms. Cell 147, 275–292 (2011).

41. Weiswald, L.-B., Bellet, D. & Dangles-Marie, V. Spherical Cancer Models in Tumor Biology. Neoplasia 17, 1–15 (2015).

42. Eilenberger, C., Rothbauer, M., Ehmoser, E.-K., Ertl, P. & Küpcü, S. Effect of Spheroidal Age on Sorafenib Diffusivity and Toxicity in a 3D HepG2 Spheroid Model. Sci. Rep. 9, 4863 (2019).

43. Pedraza, V. et al. Gene expression signatures in breast cancer distinguish phenotype characteristics, histologic subtypes, and tumor invasiveness. Cancer 116, 486–496 (2010).

44. Hanahan, D. & Weinberg, R.A. Hallmarks of cancer: the next generation. Cell 144, 646–674 (2011).

45. Sanchez-Vega, F. et al. Oncogenic Signaling Pathways in The Cancer Genome Atlas. Cell 173, 321–337.e310 (2018).

46. Biswas, D.K. et al. NF-κB activation in human breast cancer specimens and its role in cell proliferation and apoptosis. Proc. Natl. Acad. Sci. U S A 101, 10137–10142 (2004).

47. Sachs, N. et al. A Living Biobank of Breast Cancer Organoids Captures Disease Heterogeneity. Cell 172, 373–386.e310 (2018).

48. Al-Hajj, M., Wicha, M.S., Benito-Hernandez, A., Morrison, S.J. & Clarke, M.F. Prospective identification of tumorigenic breast cancer cells. Proc. Natl. Acad. Sci. U S A 100, 3983–3988 (2003).

49. Urbach, D., Lupien, M., Karagas, M.R. & Moore, J.H. Cancer heterogeneity: origins and implications for genetic association studies. Trends Genet. 28, 538–543 (2012).

50. Meacham, C.E. & Morrison, S.J. Tumour heterogeneity and cancer cell plasticity. Nature 501, 328–337 (2013).

51. Hernandez-Gordillo, V. et al. Fully synthetic matrices for in vitro culture of primary human intestinal enteroids and endometrial organoids. Biomaterials 254, 120125 (2020).

52. Magno, V., Meinhardt, A. & Werner, C. Polymer Hydrogels to Guide Organotypic and Organoid Cultures. Adv. Funct. Mater. 30, 2000097 (2020).

53. Hedegaard, C.L. et al. Peptide-protein coassembling matrices as a biomimetic 3D model of ovarian cancer. Sci. Adv. 6, eabb3298 (2020).

54. Kelm, J.M., Timmins, N.E., Brown, C.J., Fussenegger, M. & Nielsen, L.K. Method for generation of homogeneous multicellular tumor spheroids applicable to a wide variety of cell types. Biotechnol. Bioeng. 83, 173–180 (2003).

55. Sikavitsas, V.I., Bancroft, G.N. & Mikos, A.G. Formation of three-dimensional cell/polymer constructs for bone tissue engineering in a spinner flask and a rotating wall vessel bioreactor. J. Biomed. Mater. Res. 62, 136–148 (2002).

56. Phan, N. et al. A simple high-throughput approach identifies actionable drug sensitivities in patient-derived tumor organoids. Commun. Biol. 2, 78–78 (2019).

57. Sant, S. & Johnston, P.A. The production of 3D tumor spheroids for cancer drug discovery. Drug Discov. Today Technol. 23, 27–36 (2017).

58. Gras, S.L. et al. Functionalised amyloid fibrils for roles in cell adhesion. Biomaterials 29, 1553–1562 (2008).

59. Wei, G. et al. Self-assembling peptide and protein amyloids: from structure to tailored function in nanotechnology. Chem. Soc. Rev. 46, 4661–4708 (2017).

60. Caliari, S.R. & Burdick, J.A. A practical guide to hydrogels for cell culture. Nat. Methods 13, 405–414 (2016).

61. Wolf, K. et al. Physical limits of cell migration: control by ECM space and nuclear deformation and tuning by proteolysis and traction force. J. Cell Biol. 201, 1069–1084 (2013).

62. Zanoni, M. et al. 3D tumor spheroid models for in vitro therapeutic screening: a systematic approach to enhance the biological relevance of data obtained. Sci. Rep. 6, 19103–19103 (2016).

63. Minchinton, A.I. & Tannock, I.F. Drug penetration in solid tumours. Nat. Rev. Cancer 6, 583–592 (2006).

64. Meneghel, J., Kilbride, P. & Morris, G.J. Cryopreservation as a Key Element in the Successful Delivery of Cell-Based Therapies-A Review. Front. Med. (Lausanne) 7, 592242–592242 (2020).

