## Supplementary Information for "Amyloid fibril-based hydrogels for high-throughput tumor spheroid modeling"

**Materials and Methods**

**Supplementary Figures and legends**

**Supplementary Tables**

**Materials and Methods**

**General chemicals and reagents**

All the general reagents and chemicals used for the experiments were procured either from Merck (Darmstadt, Germany) or Sigma-Aldrich (St. Louis, MO, United States) and were of the highest quality. The chemicals used for cell culture and related assays are described in the respective sections. Double-distilled, deionized water was obtained using a Milli-Q system (Millipore Corp., Bedford, MA, USA) and were autoclaved.

**Designing of amyloid-based hydrogel**

The peptide-forming hydrogel was designed based on the self-assembly of high amyloidogenic segments from disease and functional amyloid protein/peptide^1-5^. Figure 1a demonstrates the design scheme for the amyloid based hydrogel peptide sequences. In general, the segment showing the highest β-aggregating regions by the aggregation-predicting algorithm TANGO^6^ was selected. These amyloidogenic peptide sequences along with Fmoc protection at the N-terminus develop the amyloid fibril network by rapid intermolecular hydrogen bond formation, which subsequently form a hydrogel by encapsulating water molecules inside a three-dimensional fibril network. Peptides A2, A8, and P5 were previously characterized and used for hydrogels in stem cell differentiation^1-3^.

**Peptide synthesis by solid-phase and ESI-Mass spectrometry**

All peptides were synthesized by a solid-phase peptide synthesis protocol using 9-fluorenylmethoxy-carbonyl (Fmoc) chemistry^7^.The synthesis was performed with a synthesis scale between ~0.20 to 0.25 mmol based on resin loading capacity i.e. ~0.25 mmol. The coupling of amino acids was done by binding of first amino acid to the resin using 0.1 eq. of 4-Dimethylaminopyridine (DMAP), 1 eq. of Fmoc protected amino acid and 1 eq. of N, N′-Diisopropylcarbodiimide (DIC) and Hydroxybenzotriazole (HOBt) in Dimethylformamide (DMF). The reaction was incubated at room temperature for 3-4 h, followed by removal of Fmoc using 20-25 % of piperidine in DMF. Finally, the free carboxyl-terminal was cleaved from the resin using a cleavage mixture of 88:5:5:2 of trifluoroacetic acid (TFA): water: phenol: triisopropylsilane (TIPS). The peptides were further precipitated out using ice-cold methyl tert-butyl ether (MTBE) with N-terminal Fmoc protected peptide. Electrospray ionization mass spectrometry (ESI-MS) experiments were performed for examining the molecular weight of peptide mass using an Agilent 6560 DTIMSQ-TOF spectrometer (Agilent Technologies, Santa Clara, CA), with the dual-ESI source operated in positive ion mode. Peptides solutions (5 mg/ml) were prepared in HPLC grade acetonitrile. The trapping funnel was tuned to avoid energizing of the complexes (RF lower than 200 V for the ion funnel, and low extraction potentials). The data were analyzed using the Agilent MassHunter software (version B.07).

**Preparation of hydrogels**

Preparation of peptide hydrogels was performed according to the previously established protocol^1-3^. In brief, for temperature-responsive gels, 1 mg of the synthesized peptide was dissolved in 200 μl of 20 mM phosphate buffer pH 7.4 by heating. Subsequently, three successive heating/cooling cycles and the addition of 150 mM NaCl were done, which lead to self-sustaining hydrogels. For pH-responsive hydrogels, 1 mg of the peptide was dissolved in 200 μl of 20 mM phosphate buffer at pH 7.4. Subsequently, the pH was increased to 10 using 2N NaOH to dissolve the peptides and then decreased and adjusted to pH 7.4 for gelation.

### MTT (3-[4,5-dimethylthiazol-2-yl]-2,5 diphenyl tetrazolium bromide) assay

### In 96-well plates (Costar, Corning, NY, USA), 30 μl of hydrogel (5 mg/ml) were added and were sterilized under UV for 30 minutes. MCF7 (~10,000) cells in 100 μl complete medium were seeded in each well and incubated for 24 h. Untreated and Triton-X (10 μl) treated cells were used as controls. The cells were further incubated for 4 h after this incubation and then 10 μl of MTT (5 mg/ml stock) were added. After this, 100 μl of 50% N, N-dimethylformamide, and 20% sodium dodecyl sulfate solution was added and incubated overnight. The absorption was recorded at 560 nm and background scattering at 690 nm was measured in a SpectraMax M2® (Molecular devices, USA) plate reader. Background scattering values were subtracted from the absorbance at 560 nm to calculate the viability of cells. viability of cells (%) = (Abs_sample_ – Abs_blank_/ Abs_control_ – Abs _blank_) x 100. The statistical significance was determined by one-way ANOVA Dunnett’s multiple comparison post-hoc test; p-values in the figure indicate *≤0.05, ** ≤0.01, ***≤0.001.

### Cryo-scanning electron microscopy

### The hydrogels were cast onto the metallic stub coated with carbon tape. The samples were frozen using a slush of liquid nitrogen (−210°C) and transferred to the preparatory chamber. The gels were fractured using an actively cooled knife followed by sublimation at −80°C for 10 min. Then sputter coated with platinum for 45 s at 10 mAmp current. JSM-7600F SEM with PP3000T automated cryo-unit (Quorum) was used for image acquisition.

### Transmission electron microscopy

### Hydrogels (5 mg/ml) were diluted (1:50 dilution in 20 mM phosphate buffer pH 7.4) and spotted individually on glow-discharged, carbon-coated Formvar grid (Electron Microscopy Sciences, Fort Washington, PA) followed by incubation for 5 mins. Subsequently, the grids were washed with distilled water followed by staining with 1% (w/v) aqueous uranyl formate solution. These grids were then observed under TEM. The image was acquired at 120 kV with magnifications in the range of 26,000X and 43,000X using a transmission electron microscope (TECNAI12 D312 FEI, Netherlands).

### Rheological measurements

### The preformed hydrogel (300 μl) was placed between a parallel plate (PP25 geometry). The dynamic oscillatory mode measurements were taken with a gap size of 0.2 mm and a constant amplitude of 0.05% at 25°C. The frequency sweep experiment was performed with an angular frequency (ω) of 0.1-100 1s^−1^ for 15 min. The step-strain oscillatory rheology with an identical setup was used to determine the thixotropicity. A high strain (100%) was applied for disruption of gel and was subsequently allowed to recover by decreasing the strain to 0.05%. The storage modulus (G’) and the loss modulus (G”) were determined and plotted as a function of time over three decades. The rheological measurements were done using Anton Paar rheometer (Graz, Austria).

### Fourier transform infrared spectroscopy (FTIR)

### Ten microliters of vortexed gel samples (5 mg/ml) were spotted on a KBr pellet that was made before the experiment by compressing ground KBr powder. Ten microliters of 20 mM phosphate buffer, pH 7.4 was spotted on other KBr pellet and utilized for the background spectrum. All samples were dried under an IR lamp. FTIR spectra were acquired by Vertex 80 FTIR system equipped with a DTGS detector (Bruker, Ettlingen, Germany). The spectra were obtained in the range of 1800–1500 cm^−1^ as an average of 32 scans, and raw data corresponding to the amide-I region (1600–1700 cm^-1^) was deconvoluted by the Fourier self-deconvolution method. The deconvoluted spectra were then subjected to a Lorentzian curve-fitting procedure using Opus-65 software. Two independent experiments were performed for each sample.

### Thioflavin T (ThT)-binding assay

### Two microlitres of a 1 mM ThT stock solution was added to 150 μl of vortexed gel (10 times diluted in the 20 mM phosphate buffer, pH 7.4), and then the solution was mixed thoroughly in a quartz cuvette. Subsequently, the fluorescence was measured immediately with a Jasco (Kyoto, Japan) using excitation at 450 nm, emission at 460–600 nm, and excitation and an emission slit width of 5 nm. ThT control experiment was done with 2 μl of 1 mM ThT dye solution mixed in 150 μl of phosphate buffer (20 mM, pH 7.4).

### Congo red (CR)-binding assay

### Fifteen microliters of CR stock solution (360 μM stock, in PBS containing 10 % ethanol) was mixed with 85 μl of vortexed hydrogel (5 times diluted) in 20 mM phosphate buffer, pH 7.4, and incubated in the dark for 5 min. Ultraviolet (UV) absorption spectra were measured from 300 to 700 nm on a UV spectrophotometer (JASCO V-650, Tokyo, Japan). CR control experiment was carried out by incubating the dye solution with the phosphate buffer (20 mM, pH 7.4).

**Cell lines and culture conditions**

MCF7, MDA MB 231 (human breast cancer cells) and HepG2 (human liver cancer cells) HeLa (cervical cancer cells), MCF 10A (human breast epithelial cells) and A549 (lung carcinoma cells) were procured from the cell repository at the National Centre for Cell Science, Pune, India. All cells are grown in tissue culture flasks under standard conditions (37°C, 5% CO_2_, 95% humidity) in DMEM media supplemented with 10% heat-inactivated fetal bovine serum. DMEM growth medium (Gibco, USA) supplemented with 2% horse serum (Gibco, USA), 0.5 μg/ml hydrocortisone (Himedia, India), 10 μg/ml insulin (Himedia, India) and 1X Pen-Strep antimicrobial agent (Himedia, India) was used for MCF 10A culture.

**Quantification of cell shape and cell area**

For evaluation of cell adhesion, 20 µl of hydrogels were placed in 24 well plates, where wells without hydrogel were treated as controls. Subsequently, 10,000 cells were added to the wells and covered with fresh medium followed by incubation at 37^°^C in air humidified atmosphere at 5% CO_2_ for 24 h. After incubation, wells were gently rinsed with PBS to remove unattached cells. Finally, the wells were imaged in a bright field using Leica Dmi8 microscope fitted with an sCMOS camera (Andor Technologies) at 20X magnification. ImageJ software was used to determine cell size and shape calculation. Cell morphology (spread area and circularity) was estimated by analyzing 5 random a frame of 50 cells/condition from three independent experiments.

**Immunocytochemistry**

To study F-actin fiber formation of cells adhered to amyloid hydrogels, immunocytochemistry was performed. Cell adhered to the tissue culture plates were served as a control. For this, MCF7 cells (~10,000) were cultured on FA1 amyloid hydrogels coated coverslip placed in 24 well plates and incubated for 24 h. After incubation, cells were washed with phosphate-buffered saline (PBS). Subsequently, cells were washed with PBS and were incubated with Phalloidin (Molecular Probes; 1:300 dilution) for observing F-actin for 1 h at room temperature. Afterward, samples were washed with PBS and stained with DAPI for nuclear staining (Molecular Probes; Invitrogen). Lastly, the coverslips were mounted and on the glass slides and were observed under Leica Dmi8 at 20X magnification in fluorescent mode. The distributions of focal adhesions/F-actin were analyzed using ImageJ. Three independent experiments were performed for this assay.

### Cell proliferation assay

For the *in vitro* cell proliferation studies, 20 µl of the amyloid hydrogels were placed in a 96-well plate followed by 1-hour UV light sterilization. Afterward, approximately MCF7 cells (~10,000) were seeded in the wells and incubated for 48 h at 37^0^C incubator with a humidified atmosphere of 5% CO2. Non-treated wells (without hydrogel) were used as a negative control. Matrigel served as a positive control to study cell proliferation. After this incubation, 10 μl MTT from (5 mg/ml stock) was added to the wells and the cells were further incubated for 4 h. Lastly, 100 μl of solubilization solution containing 50% N, N-dimethylformamide, and 20% sodium dodecyl sulfate solution was added, and the mixture was incubated overnight. The absorption was recorded at 560 nm using a SpectraMax M2e microplate reader (Molecular Devices, USA). The absorbance reading of the blank (only medium) was subtracted from all samples. Absorbance readings from test samples were then divided by corresponding controls and multiplied by 100 to give a percentage of cell viability or proliferation. Absorbance values greater than the control indicate cell proliferation, while lower values suggest cell death or inhibition of proliferation. Three independent experiments were performed for this assay.

**Wound healing assay**

To determine the effect of amyloid hydrogel on cell motility (migration capability) wound healing assay was performed^8^. MCF7 cells (~50,000) were cultured on the 6 well plates at 37°C. Once the confluence of MCF 7 monolayer cells was achieved, the wells were scratched with a 200 ul sterile pipette tip to create a 0.25 mm size mimicking an incision wound. Then, the plates were washed with PBS to remove cell debris. After that, the wound was covered with PBS as control, each amyloid hydrogel with bFGF (2 µM), and incubated for 5 mins to solidify the hydrogel. The wells were covered with 1 ml of fresh cell culture medium and incubated at 37°C, 5% CO_2_ in an air humidified incubator. Subsequently, the closure of the wound was imaged at 0, 12, 24, 36, 48, and 72 h to investigate and analyze the scratch wound assay using different samples. The scratch area was measured using the Image-J software. The images were captured using Leica Dmi8 10X magnifications fitted with an sCMOS camera (Andor Technologies, USA). Wound width was estimated using the ImageJ software. Cell migration was calculated using the formula: Wound size (%)=100 × [initial wound width - final wound width)/ initial wound width)]. n=3 independent experiments were performed.

**Three-dimensional cell culture using amyloid hydrogels**

First, 20 µl of the amyloid hydrogels in 20 mM phosphate buffer, pH 7.4 were coated on a glass coverslip placed in 24 well plates followed by 30 mins UV light sterilization. Subsequently, cells were trypsinized using 1X trypsin-EDTA, and 10,000 cells were pelleted and mixed with 20 μl of vortexed hydrogel in 20 mM phosphate buffer, pH 7.4. This gel-cell mixture was dispersed throughout the hydrogel coated cover slip and incubated for 10 minutes for hydrogel to solidify. Fresh medium (DMEM media supplemented with 10% heat inactivated fetal bovine serum) was added in the wells slowly without disturbing and kept for 10 days for incubation. The formation of cell aggregates/spheroid was visualized every alternate day by capturing images using Leica Dmi8 at microscope with 10X magnifications fitted with an sCMOS camera (Andor Technologies, USA). Diameter and volume of the spheroid were quantified every alternate day for all spheroids using ImageJ software and analyzed the size of 3D cellular structures. Further, aggregation of cells or spheroid shape (roundness metrics) and solidity was estimated using Image J software. Shape represents the object's roundness (1 for a perfect circle; 1> for less spherical objects). Cell density was measured in terms of solidity, which is the average optical density (intensity) value of all the pixels of a selected object^9,10^. It means that the more compacted cells are inside the aggregates/spheroids, the more cells will be placed to the same point in the two-dimensional image and hence, the brightness will be less in each pixel of an image ^9,10^.

**Cell viability assay in 3D cell culture**

The cell viability assay in 3D cell culture was performed using Calcein-AM/Ethidium homodimer-1 staining. Following incubation in the medium for days as mentioned above for 3D cell culture, cellular aggregates/spheroids were stained with a mixture of two dyes composed of 1 μM calcein AM and 1 μM of ethidium homodimer-1 (LIVE/DEAD™ Viability/Cytotoxicity Kit, Thermo Fisher, USA)^11^. Dyes were prepared freshly in sterile phosphate-buffered saline (PBS), pH 7.4, and added directly to the media without aspiration. Since, dye penetration is slower into spheroids than 2D cultures, spheroids were incubated with dye for 2 h before imaging. The dye solution was washed out with PBS slowly, and care was taken during pipetting to avoid spheroid loss, disintegration, or displacement. After incubation, the spheroids were observed and imaged under a Leica DMi8 microscope in fluorescent mode with 20X magnification. Three independent sets of experiments were performed.

##### Immunofluorescence study

To analyze cell adhesion molecules such as cadherin and integrin, cell aggregates/spheroids in 3D culture and 2D monolayer culture were performed for 3 days. For the detection of epithelial to mesenchymal transition (EMT) markers, immunostaining was formed with 3D cultured cells and 2D monolayer culture after 3, 5, and 7^th^ days of incubation. After appropriate days of culture, cell aggregates in 3D were washed with PBS gently. Subsequently, cells were fixed in 4% paraformaldehyde for 1 h in the dark at room temperature. After fixation, cells were permeabilized using 0.5% Triton X-100 for 20 minutes, blocked with 1% BSA for non-specific epitopes for 1 h. After PBS wash, the cells in 3D culture and 2D culture were then incubated with rabbit monoclonal primary antibody against E and N-cadherin, β-integrin, Slug, and VEGF (CST) (1:500 dilutions each) at 4 °C for 16 h. AlexaFluor 555 conjugated anti-rabbit secondary antibody (Life Technologies, USA) and AlexaFluor 488 conjugated anti-rabbit secondary antibody for VEGF was used in 1:1000 dilution each for 2 h followed by washing with PBST, pH 7.4 and stained with 1 μg/ml DAPI for 5 mins and were washed twice with PBS. Further, cells were rinsed three times with PBST. Cells in 3D and 2D were then visualized under laser scanning confocal microscopy at 40X magnification (Carl Zeiss, LSM 780) and the intensity of fluorescence was analyzed using Image J software. Three independent experiments were performed.

**Western blots analysis**

Cells in 3D culture and 2D monolayer culture of MCF7 were collected after 3 days of incubation. Subsequently, cells were washed and disintegrated using 1 ml of Corning® cell recovery solution and collected with cold PBS. Whole-cell lysates were prepared using a buffer containing 50 mM Tris-HCl, pH 7.5, 250 mM NaCl, 5 mM EDTA, 0.5% NP-40, a cocktail of protease inhibitors (Roche Diagnostics). Total protein was measured using Bradford assay and an equal amount of protein was denatured with a Lamelli sample buffer containing 100 mM DTT for 10 min. Samples were resolved in 12% SDS-PAGE and proteins from the gel were transferred to a nitrocellulose membrane by wet transfer method using BioRad transfer apparatus. The transfer was performed at 4°C for 2 h at constant current (350 mA). After transfer, the membrane was washed with TBST (0.01% Tween) followed by blocking with 5% skimmed milk powder for 1 h at RT under rocking condition. Two washes (10 min each) were given to the membrane with Tris-buffered saline (50 mM Tris, 150 mM NaCl). Immunodetection on the membrane was carried out with primary rabbit monoclonal antibody; anti-E cadherin and anti-β-catenin (1:1000 dilution for each, Cell signaling technologies, USA) and mouse anti- GAPDH (1:2000; Santa Cruz Biotechnology, USA) followed by an anti-rabbit antibody and anti-mouse (1:3500 dilution each) conjugated to horseradish peroxidase (Calbiochem) for 2 h at room temperature under constant rocking. Afterward, two washes (10 min each) were given with TBST (0.01% Tween) and the blot was treated with Supersignal west pico-chemiluminescent substrate (Thermo Scientific, USA) and visualized on X-ray film (Amersham, USA). Images were processed using Image J software for quantitative analysis. Three independent experiments were carried out.

**Drop-Cast (DC) method for single tumor spheroid formation in 96-well plate**

First, 10 µl of the amyloid hydrogels in 20 mM phosphate buffer, pH 7.4 were placed in 96 well plates followed by 30 mins UV light sterilization. Subsequently, cells were trypsinized using 1X trypsin-EDTA, and 10000 cells were pelleted and mixed with 10 μl each of the hydrogels. Matrigel was used as a control. 10 μl mixture of cells and hydrogels were then seeded in a form of drop on the coverslip (coated with corresponding preformed gel) with micropipette tip followed by incubation for ten minutes for hydrogel to solidify. Fresh medium (DMEM media supplemented with 10% heat-inactivated fetal bovine serum) was added in the wells slowly without disturbing and kept for 10 days for spheroid formation. The formation of a single spheroid was visualized every alternate day using the Leica Dmi8 microscope at 10X magnifications fitted with a sCMOS camera (Andor Technologies, USA). The diameter of the spheroid was quantified every alternate day for all spheroids using ImageJ software, which assesses the size of 3D cellular structures. The viability of cells was analyzed through Trypan Blue cell exclusion assay.

Further, the cell viability was done using Calcein AM/Ethidium homodimer-1 staining. For this, spheroids were cultured for 3,5 and 7^th^ day in a drop cast method, the media was taken out and spheroids were washed twice with DPBS (Dulbecco’s PBS) to remove residual serum and samples. Drop cast spheroids were then incubated in presence of 50 μM Calcein AM and 10 μM Ethidium homodimer-1 (LIVE/DEAD™ Viability/Cytotoxicity Kit, Thermo Fisher, USA) for 30 min at room temperature in dark. Calcein AM/Ethidium homodimer-1 signal was then checked and imaged under laser scanning confocal microscopy at 10X magnifications (Carl Zeiss, LSM 780) and images were analyzed using Image J software.

**Cytotoxicity assays in the presence of various anticancer drugs**

Drop cast spheroids and corresponding 2D monolayer culture were incubated for three days in 24 well plates in DMEM (Dulbecco’s modified Eagle’s medium) with 10% FBS, in a 5% CO_2_ incubator at 37°C followed by treatment with different chemotherapeutic drugs (cisplatin, doxorubicin, paclitaxel, pentostatin, and 5-Fluorouracil; 0-14 µM) for 24 h. For control, a media containing buffer was used. After 24 h of incubation, 10 μl of MTT solution (5 mg/ml prepared in PBS) was added to each well followed by incubation for 4 h. Subsequently, 100 μl of SDS-DMF solution (50% DMF and 20% SDS, pH 4.75) were added and kept for overnight incubation. The absorption values at 560 nm and 690 nm (background absorbance) were determined using a SpectraMax M2e microplate reader (Molecular Devices, USA). The background scattering values at 690 nm were subtracted from the absorbance values at 560 nm to calculate the viability of cells both in the presence and absence of various compounds at each drug concentration. IC_50_ of each drug was calculated using GraphPad Prism 8.

### RNA isolation and quantitative real-time PCR

To investigate the expression profile of potential cancer biomarkers in tumor spheroid cells (cultured in 3D drop cast method) in comparison to 2D culture, RNA was isolated from MCF7, MDA MB 231, Hep G2, A549, HeLa spheroids, and their respective monolayer culture followed by cDNA synthesis. Gene expression was analyzed by quantitative Real-time PCR. Markers analyzed have implications in cell cycle regulation, differentiation, and malignant transformation^12, 13^. RNA was isolated using TRIzol reagent (Invitrogen, Carlsbad, CA, USA) according to the manufacturer’s protocol. Briefly, the cell pellet was lysed using 700 μl of TriZol. Further, 400 μl of chloroform was added and samples were vortexed vigorously. Samples were centrifuged at 12,000 g for 15 mins at 4°C. The aqueous phase (upper layer) was transferred to fresh tubes and an equal volume of isopropyl alcohol was added for precipitation of RNA. Samples were incubated at room temperature for 20 mins and centrifuged at 12,000 g for 10 mins at 4°C. The supernatant was discarded and the RNA pellet was washed using 70% ethanol followed by centrifugation at 7,500 g for 5 mins at 4°C. Ethanol was allowed to evaporate and RNA was resuspended in nuclease-free water. The concentration of the isolated RNA was measured in a nanodrop spectrophotometer (Implen, USA). Total RNA was then reverse transcribed to cDNA with a ProtoScript First Strand cDNA Synthesis Kit (NEB, Ipswich, MA, USA) using random hexamers and Oligo(dT) 20 primers according to the manufacturer’s protocol. Real-time PCR (qRT-PCR) was performed using the SYBR Green method using primers for genes (Supplementary Table 1 for primer sequences). Predesigned and validated SYBR green primers were purchased from Sigma-Aldrich (Bangalore, India) for the study. Maxima SYBR Green/ROX qPCR Master Mix (2X) (ThermoFisher Scientific, USA) was used according to the manufacturer’s protocol. n=3 independent experiments were performed.

**Microarray hybridization**

For microarray experiments, spheroids were cultured and RNA extraction was done as described above. The concentration and purity of the RNA were evaluated using the Nanodrop Spectrophotometer (Thermo Scientific; 2000). The integrity of the RNA was analyzed on the Bioanalyzer (Agilent; 2100 expert). We considered RNA to be of good quality based on the 260/280 values (Nanodrop) and optimal RNA integrity profile in Bioanalyzer**.** The microarray hybridization and scanning were performed at the Agilent certified microarray facility of Genotypic Technology, Bengaluru, India. Gene expression samples were labeled using the Agilent Quick-Amp labeling Kit (p/n5190-0442). Reverse transcription of total RNA was done using oligo dT primer at 40°C tagged to a T7 polymerase promoter and to make double-stranded cDNA. Further, for cRNA generation, synthesized double-stranded cDNA was used as a template followed by incorporation of Cy3 CTP(Agilent) dye at 40°C. Subsequently, using Qiagen RNeasy columns (Cat No: 74106, Qiagen) labeled cRNA was cleaned up followed by the assessment of yield and specific activity using the Nanodrop ND-1000. Fragmentation of Labeled cRNA samples was done using the Gene Expression Hybridization kit of Agilent Technologies (Part Number 5190-0404, In situ Hybridization kit,) at 60ºC and hybridized on to an Agilent Human Gene Expression Microarray 8 x 60K. Hybridization was carried out in Agilent’s Surehyb Chambers at 65º C for 16 hours. The slides were washed using Agilent Gene Expression wash buffers (Part Number 5188-5327, Agilent Technologies) and scanned using the Agilent Microarray Scanner (Part Number G2600D, Agilent Technologies).

**Microarray Data Analysis**

Raw data were extracted from images using Agilent Feature Extraction software and analyzed using Agilent GeneSpring GX (v14.5) software. Further, data normalization was done using the 75^th^ percentile shift method in GeneSpring GX [Percentile shift normalization is a global normalization, where the locations of all the spot intensities in an array are adjusted]. For this normalization, each column in an experiment was taken independently, and the percentile of the expression values was calculated across all spots (where n ranges from 0-100 and n=75 is the median). This expression value is subtracted from each entity and fold change values were estimated by comparison of test samples with respect to a specific control. Significantly upregulated genes with fold change >=1 (logbase2) and downregulated genes with fold change <=-1 (logbase2) in the test samples with respect to control were obtained. Statistical analysis using student t-test and respective p-values among the replicates was estimated based on the Volcano Plot Algorithm. Differentially regulated genes were clustered using hierarchical clustering based on the Euclidean distance method and complete linkage with row and column dendrograms and plotted by Bio Vinci program (Bioturing Inc, San Diego). The biological analysis was done for the differentially expressed genes based on their functional category and pathways using the Biological Analysis tool DAVID (<http://david.abcc.ncifcrf.gov/>)^14^. Annotation for biological process, cellular components, and molecular function were analyzed by STRING and plotted by GraphPad prism software. Gene enrichment, oncogenic signatures, and pathway enrichment were analyzed by Metascape analysis <https://metascape.org/gp/index.html>^15^.

**Animal handling**

SCID mice of 6-8 weeks old, weighing ~20–25 g was procured from the ACTREC, Mumbai, India, and was acclimatized following transportation to the new location for a week. All animals were maintained in-house under standard conditions that are 50% humidity, a temperature of 22 ± 2°C having a 12 h dark/light cycle, kept in micro-isolator cages with autoclaved bedding. Food and water were provided ad libidum. The study (Approval certificate: ARI/IAEC/2020/13) was approved by the Institutional Animal Ethics Committee, Agharkar Research Institute, Pune.

**Mouse xenograft using MCF7 cells**

MCF7 cells (human breast cancer cells) were used for tumor induction. The mice were injected subcutaneously in the mammary fat pad with 2x10^7^ healthy MCF 7 cells (200 μl each injection) in sterile PBS (150 μl each injection) and only PBS was injected as a control. Subsequently, Estradiol (Estrabet tablets, 2 mg, Abbott India Ltd, India) was added to their drinking water (1000 nM i.e. 0.054 µg/25 g mouse/day). To observe the tumor formation, the mice were weighed once a week and the area of injection was monitored. At the end of the experiments, the tumors were excised and stored in cold PBS followed by processing for single-cell isolation. The experiment was repeated twice with mice in groups of 4, each time.

**Tumor tissue processing**

Tumor tissue was processed according to the protocol published recently^16^. In brief, tumor tissues were snapped after arrival and chopped into 1-3 mm^3^ pieces. Four random pieces were snapped frozen and stored at −80°C for DNA isolation, four random pieces were fixed in formalin for histopathological analysis and immunohistochemistry, and the rest was processed for the isolation of viable cancer cells. For this, the tissue was minced, washed with (Advanced DMEM/F12 containing 1X Glutamax, 10 mM HEPES, and antibiotics) and digested in 10 mL spheroid medium (Table 3) containing 1-2 mg·ml^-1^ collagenase on an orbital shaker at 37°C for 1 h. The suspension was given a short spin to remove large chunks of tissue pieces. Subsequently, 2% fetal calf serum (FCS) were added to the suspension before centrifugation at 1200 rpm. The pellet was resuspended in 10 ml of spheroid medium. Cells were counted using a hemocytometer. Viability of cells was observed by trypan blue.

**3D cell culture and drop cast spheroid using cells from tumor tissues.**

***3D cell culture:*** 20 µl of each of the hydrogels (FA1 and FA4) were spread on a 10 mm coverslip as a bed for forming spheroids. Subsequently, 20 µl from 5 mg/ml of hydrogel (FA1, FA4, and Matrigel) were mixed with 10,000 cells and spread over the coverslip and incubated at 37°C, 5% CO2, 95% humidity in tumor spheroid media (Supplementary Table 3) for 7 days. At regular intervals, cellular aggregation/spheroid formation was imaged in phase contrast microscope using Leica Dmi8 fitted with a sCMOS camera (Andor Technologies) with 10X magnifications.

***Drop cast method:*** 20 µl of each of the hydrogels (FA1 and FA4) in 20 mM phosphate buffer, pH 7.4 was spread on 10 mm coverslip as a bed for forming spheroids. Subsequently, 10 µl of hydrogel was mixed with 10,000 cells and drop cast in the form of drop on a coverslip and was allowed to solidify at 37°C for 20 min. Upon complete [gelation](https://www.sciencedirect.com/topics/immunology-and-microbiology/gelation), 400 μL of tumor spheroid medium (Supplementary Table 3) was added to each well, and plates were transferred to humidified 37°C / 5% CO_2_ incubators. Formations of spheroid were imaged in phase contrast microscope of Leica Dmi8 fitted with a sCMOS camera (Andor Technologies) with 10X magnifications. The medium was changed every 4 days and spheroid growth was monitored. Cell viability was using Trypan Blue cell exclusion assay.

**Flow cytometry analysis**

For relative quantification of cell death and apoptosis in the presence of cisplatin, 5-fluorouracil, paclitaxel, and doxorubicin drugs, flow cytometry was performed using Annexin V-FITC apoptosis detection kit (Sigma, USA). To do that, drop cast spheroids were cultured for 3 days (~10^6^ cells). On the 3rd day of spheroid culture, fresh media containing 4, 6, 8 and 12 µm of cisplatin, 5-fluorouracil, paclitaxel doxorubicin, and the buffer was added and the plate was incubated for 24 hrs. After incubation, the cells were trypsinized, centrifuged, and used for cell death assay using Annexin V-FITC Apoptosis detection kit (APOAF, Sigma, USA). The cell pellet was washed with 1X PBS and subsequently resuspended in 1X binding buffer (Sigma, USA). Cells were stained with Annexin V-FITC and propidium iodide (PI) according to the manufacturer's instructions. Unstained cells (without Annexin V-FITC and PI) were used as a control and cells stained with either Annexin V-FITC or PI were used as fluorescent compensation controls. AnnexinV-FITC and PI staining were quantified using a flow cytometer (FACSAria, BD Biosciences, San Jose, CA) and analyzed using the BD-FACS Diva software. For each sample, 20,000 cells were analyzed.

**Sorting of CD44^+^/CD24^-^ cell population in tumor spheroids and isolated tumor from mice xenograft model**

Tumor spheroids reconstructed by drop cast method using cells of mice xenograft tumors using FA1 and FA4 hydrogels were cultured for 3 and 5 days in tumor spheroid culture medium in 24 well plates. Dissociated single cells from dissected tumors were served an *in vivo* tumor control. Spheroids were disintegrated into single cells by Corning® Cell Recovery solution. Disintegrated cells and in vivo control samples were stained for 90 min on ice in tumor spheroid media with the following antibodies: PE-conjugated anti-CD24 (Biolegend, San Diego, CA), and FITC conjugated anti-CD44 [1:100] (clone IM7, Biolegend #103039). After incubation, cells were pelleted, rinsed, and resuspended in culture media and analyzed by FACS Aria (BD Biosciences, San Jose, CA) and BD FACS Diva software. Dead cells and debris were first excluded based on size via a bivariate plot of forward scatter (FSC) vs. side scatter (SSC). Cells events were analyzed for CD24 and CD44 expression on a bivariate plot. Four cell populations: CD24^−^/CD44^−^, CD24^+^/CD44−, CD24^−^/CD44+ and CD24^+^/CD44^+^ were obtained. Data were analyzed and plotted with the help of FlowJo™ Software.

**Cryopreservation of *in vitro* generated tumor spheroid**

Spheroids from mice xenografted tumor cells were cultured for three days on FA1 gel in tumor spheroid media in duplicate 35 mm plates. One plate was tested for cell viability and stained with (LIVE/DEAD™ Viability/Cytotoxicity Kit) according to the procedure mentioned above. 200 μL of cryopreservation solution was added carefully to the spheroids in the plates (60:30:10 ratio of FBS: media: DMSO) for cryopreservation. This plate was placed in liquid nitrogen and stored for two months. To thaw the spheroids, 1 mL of warmed cell culture medium was added to the 35 mm plate. Next, the medium was removed, and the spheroids were washed with cell culture medium. Finally, the viability of spheroids was analyzed by (LIVE/DEAD™ Viability/Cytotoxicity Kit) according to the procedure mentioned above and imaged using Leica Dmi8, microscope fitted with a sCMOS camera (Andor Technologies) at 10X magnifications in a fluorescent mode. Three independent experiments were performed.

**Statistical analysis**

The statistical data were expressed as mean ± s.e.m with a one-way analysis of variance (ANOVA) using the Graph Pad Prism software. The significance of differences was statistically tested with a one-way analysis of variance, followed by a Dunett’s multiple comparison post-hoc test; p-values for each graph/plot are mentioned in the corresponding figure legends

**Supplementary Figures**

**Supplementary figure 1:**

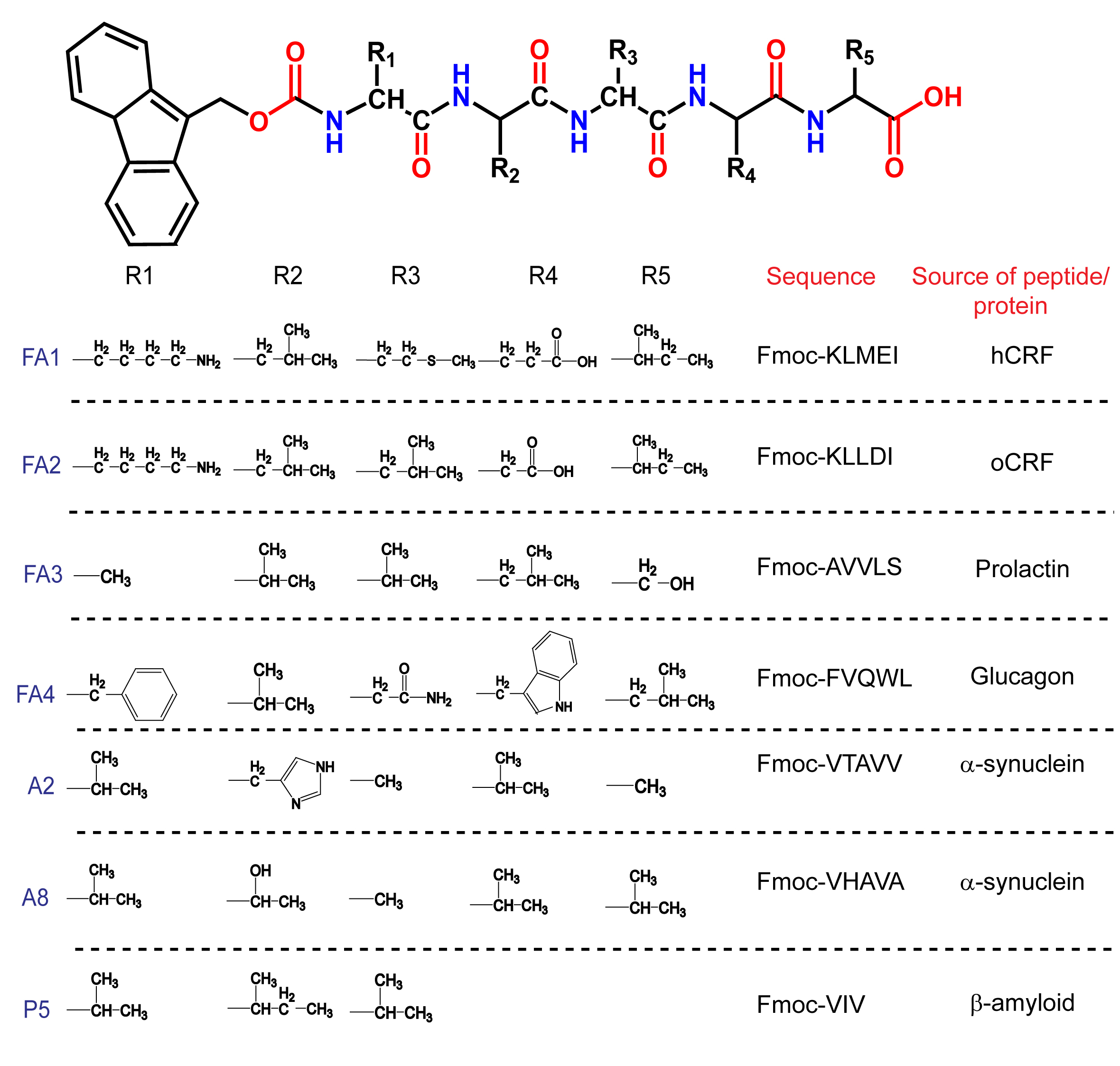

**Figure S1. Peptide sequences used for hydrogel formation.** Peptide sequences for amyloid hydrogels formation are designed from highly amyloidogenic sequences based on TANGO analysis^6^. The peptide sequences along with their corresponding source peptide/proteins are shown. A2, A8, and P5 were previously shown to form amyloid hydrogel formation for stem cell differentiation^1-3^. The N-terminus of the peptide was protected with a Fmoc group, which is known to enhance the intermolecular π- π stacking interactions.

**Supplementary figure 2:**

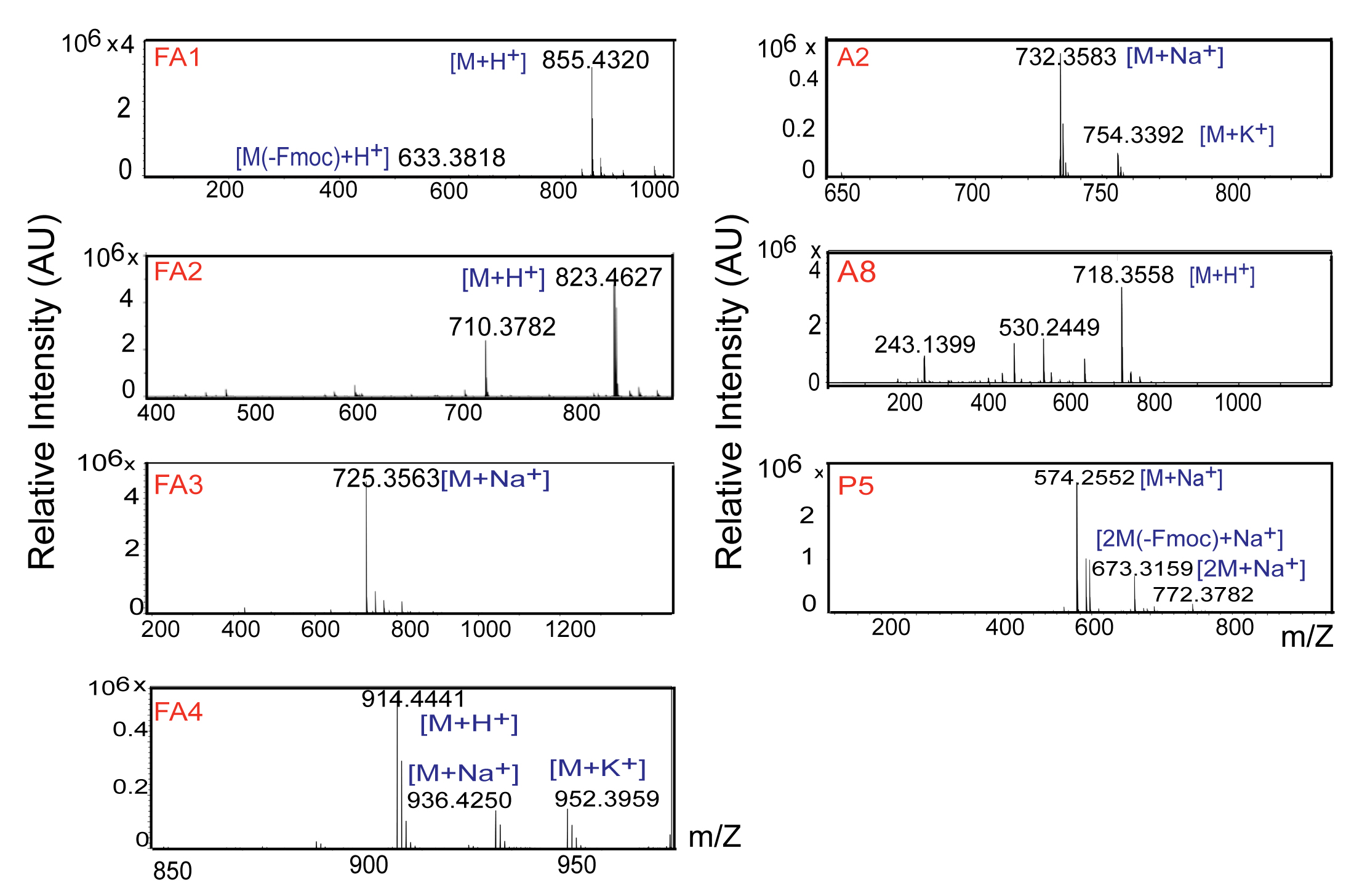

**Figure S2. ESI-Mass spectrometric analysis of synthesized peptides.** ESI-MS/MS spectrum of the synthesized peptides with m/z showing fragmentation consistent with the molecular weight of the peptide sequence. The ESI-MS spectrum of the peptide showing a major peak with the highest intensity corresponding to their molecular masses of the peptide sequence. Three independent experiments are performed.

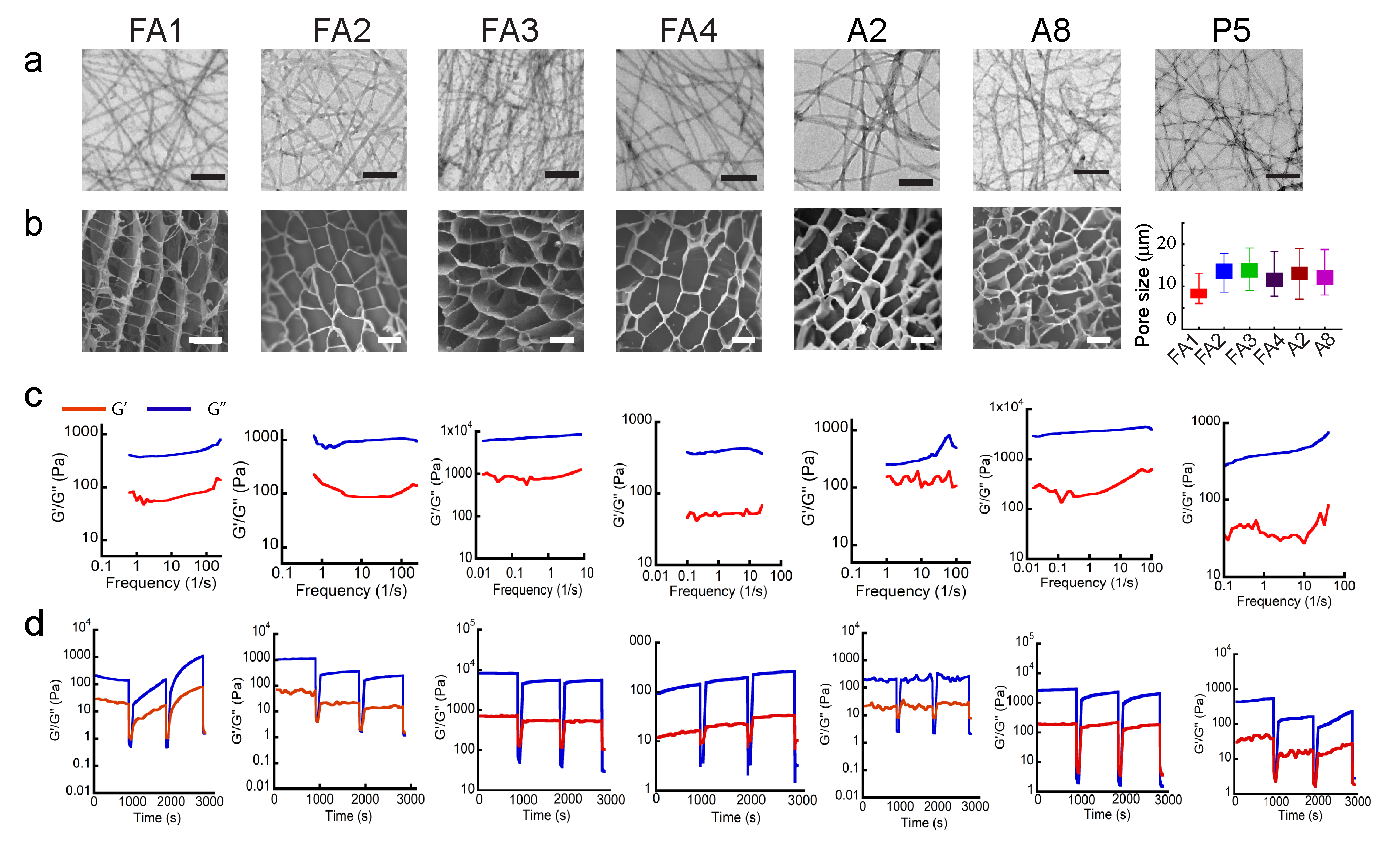
**Supplementary figure 3:**

**Figure S3. Biophysical characterization of the amyloid hydrogels**. (a) Electron microscopic images of dried hydrogel showing dense long fibrils and non‐branched filaments. The scale bar is 200 nm. (b) Morphological characterization under cryo-SEM showing that hydrogels are made of porous filamentous structure (left panel). The scale bar is 10 µm. Right panel showing pore size from SEM image analyzed by Image J indicating FA3 has the largest pore size while A8 has the smallest one. Mean pore sizes of the amyloid hydrogels range from 9-17 µm. (c) The plot of the storage modulus (G’) vs the frequency of the hydrogels is displayed. Rheological characterization revealed that at a lower frequency, the storage modulus G’ is higher than that of the loss modulus G” by approximately one order of magnitude indicating the formation of hydrogels. (d) The stress-strain rheology measurement showing the thixotropic nature of hydrogel. Three independent experiments are performed.

**Supplementary figure 4:**

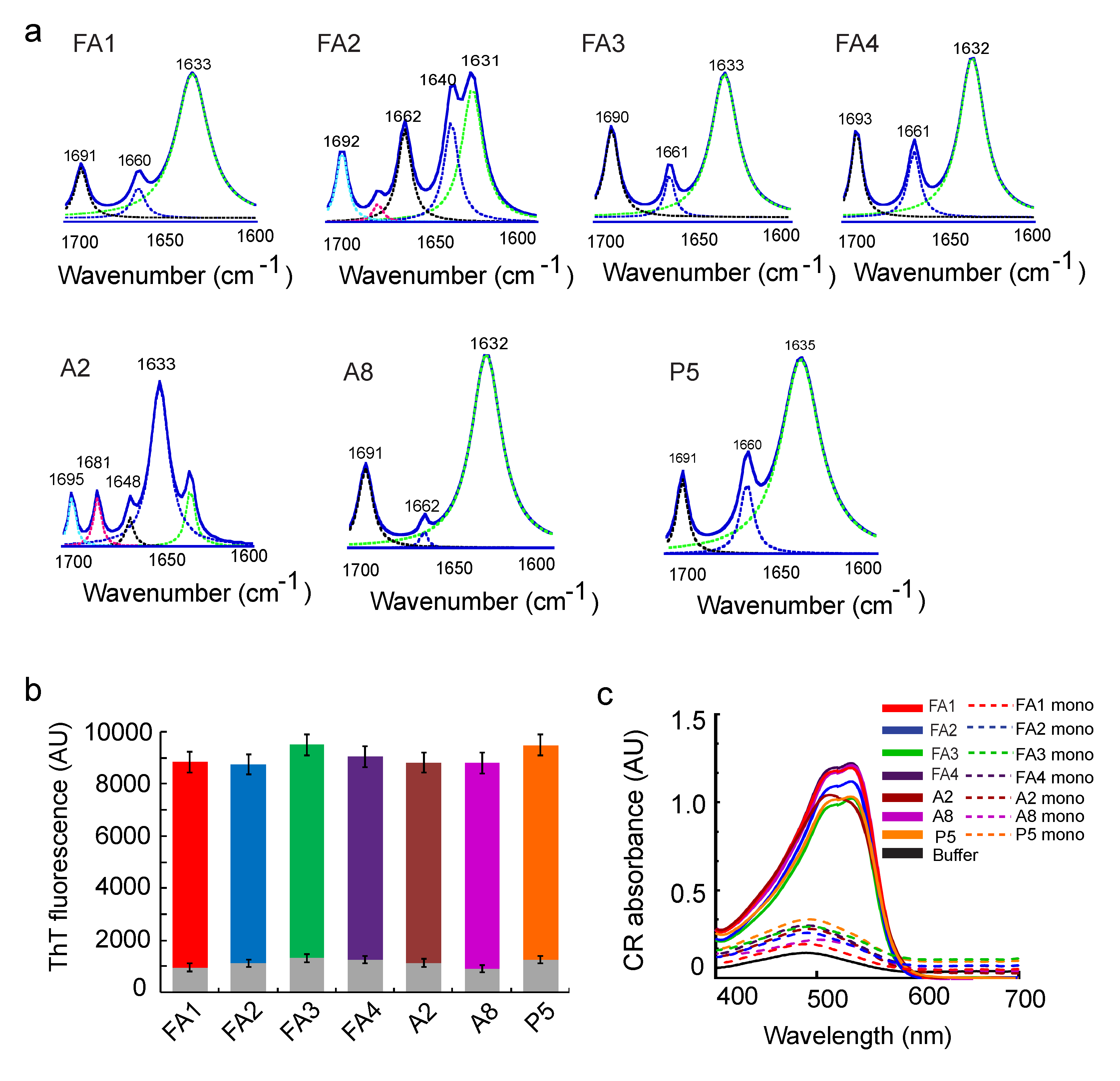

**Fig. S4. Hydrogels are composed of amyloid fibrils.** (a) FTIR of spectra of dried amyloid hydrogels indicating peaks at the amide-I region at ~ 1630 and 1690 cm^−1^ depicting typical cross β-sheet structures. (b) Thioflavin T (ThT) fluorescence of the hydrogels was done. The high ThT fluorescence intensity at 480 nm after binding to the hydrogels indicates the presence of amyloid fibrils in the hydrogels. The grey bar inside the gels represents the ThT binding of the corresponding monomeric peptides. (c) Congo red absorption plot of all amyloid hydrogels depicting a redshift and increased absorption at 520 nm indicating the amyloidogenic nature of the hydrogels. n=3 independent experiments.

**Supplementary figure 5:**

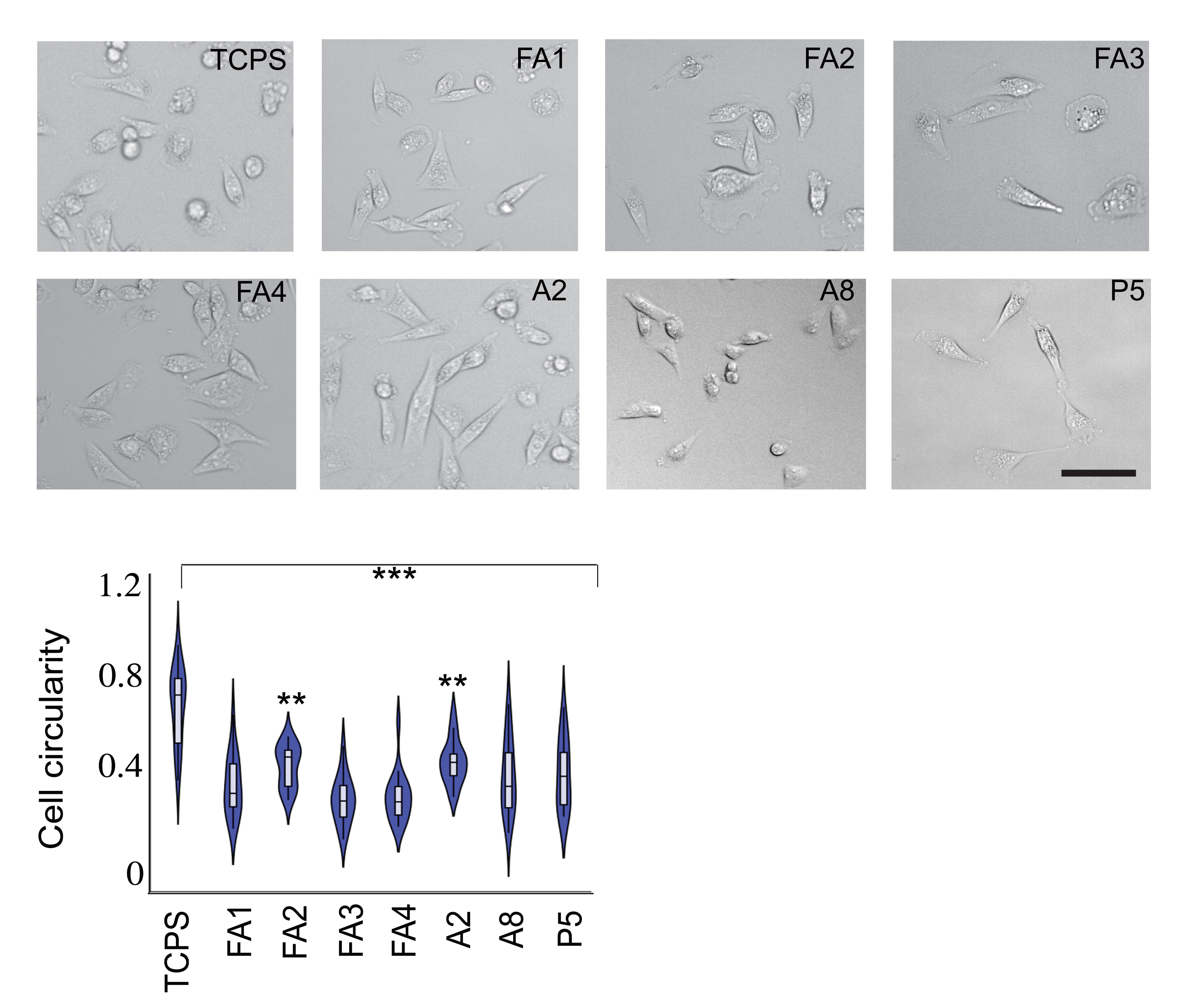

**Figure S5. Cell adhesion on the amyloid hydrogels.** Representative bright-field images indicating the morphology of MCF7 cells cultured on various amyloid hydrogels (upper panel). The scale bar is 50 µm. The circularity plot measured by ImageJ showing a decrease in circularity when cells were cultured on hydrogels (lower panel). Data plotted represent mean ± s.e.m, n=3 independent experiments. Statistical significance (***p ≤ 0.001, **p ≤ 0.01).

**Supplementary figure 6:**

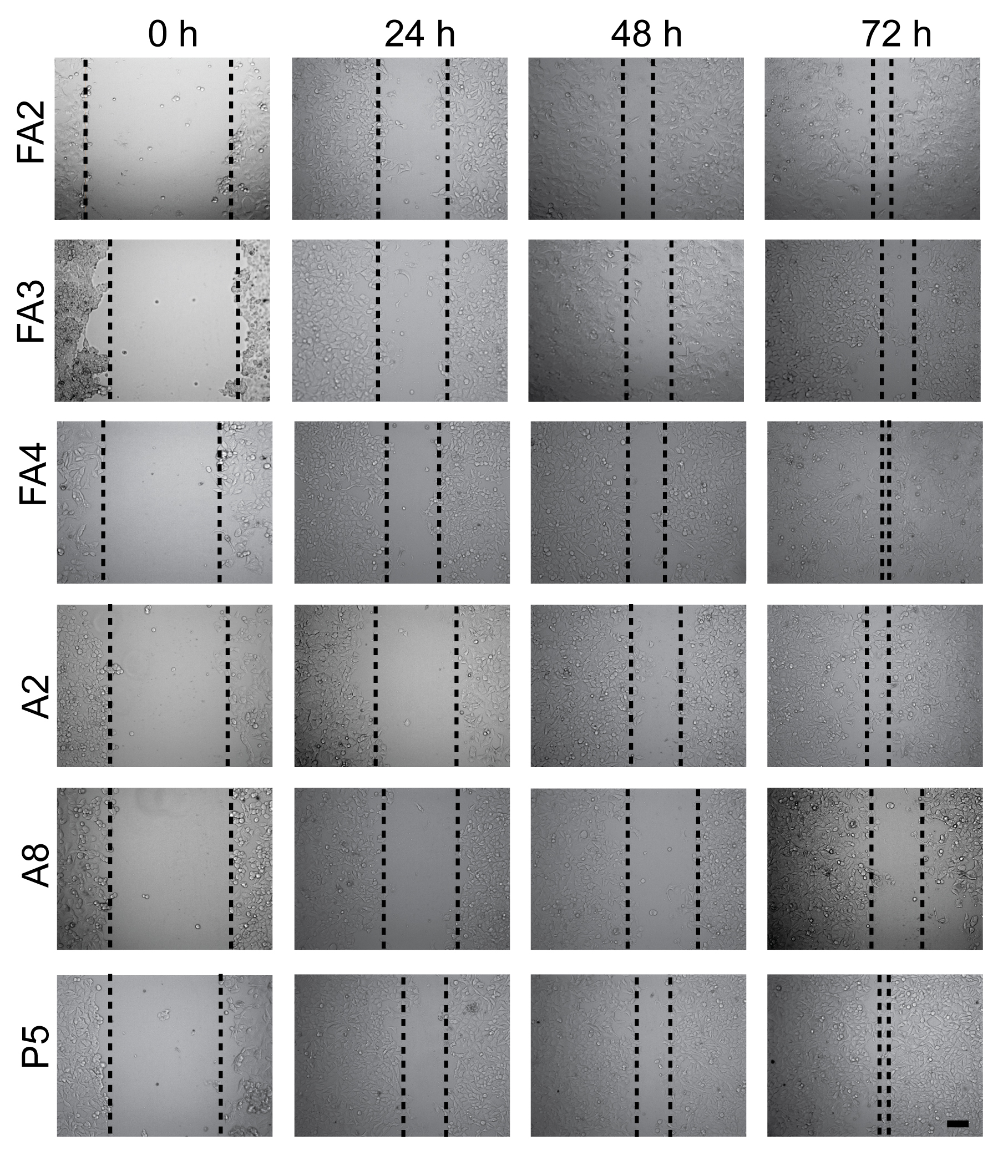

**Figure S6. Cell migration on amyloid hydrogel by wound healing assay.** Time-dependent bright-field images showing wound healing of MCF7 cells cultured on the amyloid hydrogel. Images showing wound closure rate is faster for cells cultured on amyloid hydrogel as compared to the control with time. The scratch area was measured using the Image-J software with 20X magnification. The scale bar is 100µm. n=3 independent experiments.

**Supplementary figure 7:**

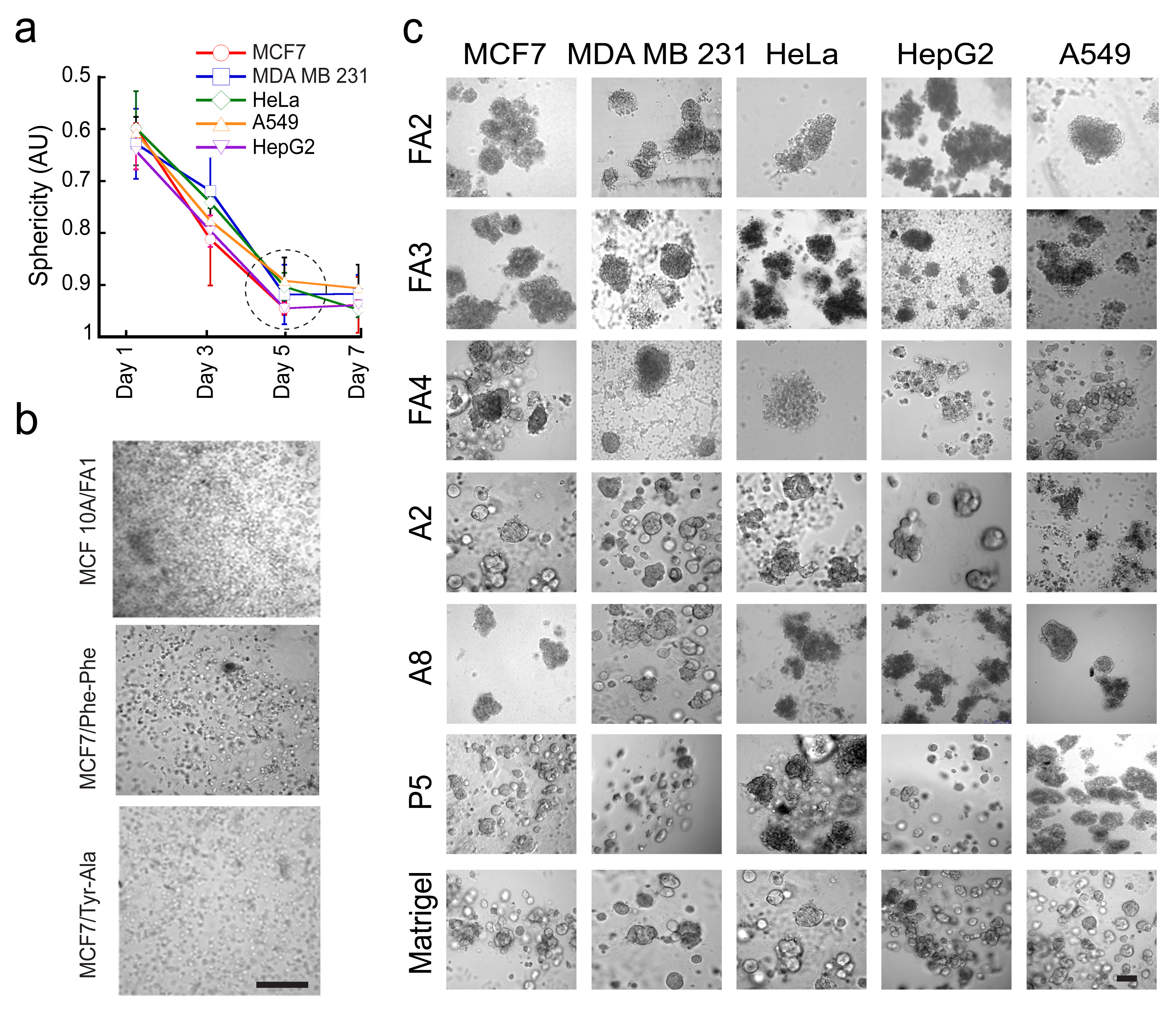

**Figure S7. Three-dimensional (3D) cell culture of MCF7, MDA MB231, HepG2, HeLa, and A549 cancer cells using amyloid hydrogels.** (a) Five different cancer cells in 3D cell culture using FA1 hydrogel showing an increase in sphericity of cellular aggregates from day 1 to day 5, which remain the same till day 7. The shape of cellular aggregates was measured with time using Image J. The quantified circularity (sphericity) was plotted against time which ranges from 0 (for a straight line) to 1 (for a perfect circle). The data represents mean ± s.e.m, n=4 independent experiments. (b) The 3D cell culture of MCF 10A, (non-tumorigenic epithelial cells, served as a negative control) in FA1 hydrogel and MCF7 cells in non-amyloid hydrogels of Fmoc-Tyr-Ala and Fmoc-Phe-Phe showing no cellular aggregation after the day 7 of incubation. The scale bar is 100 μm. (c) The 3D cell culture of various cancer cells in all amyloid hydrogels showing cellular aggregates formation on day 7. The data indicating that irrespective of the different peptide sequences, all amyloid hydrogels are capable of inducing spheroid formation for multiple cancer cell types. The scale bar is 100 µm. n=4 independent experiments.

**Supplementary figure 8:**

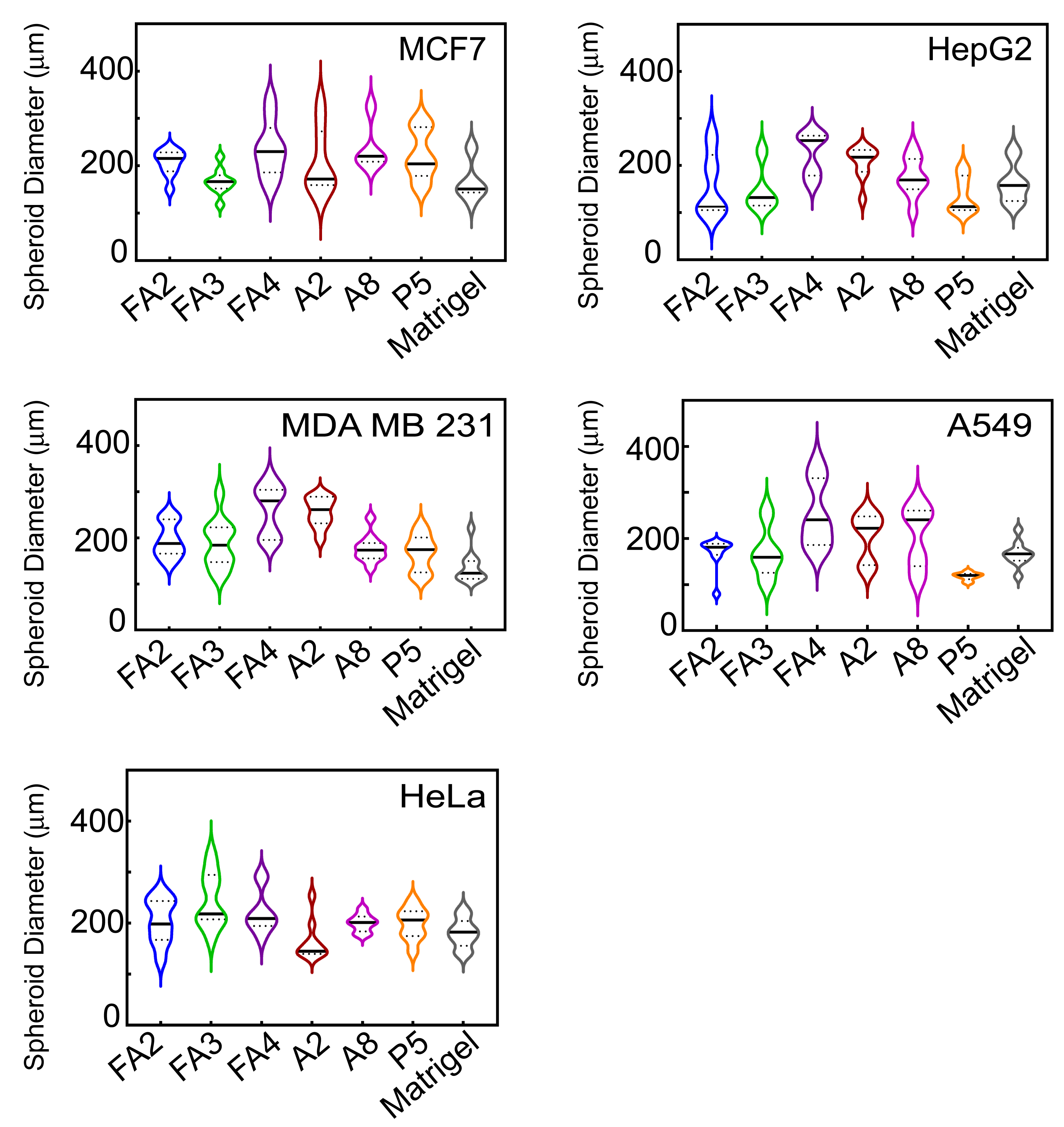

**Figure S8. The diameter of day 7 spheroids of various spheroids on different amyloid hydrogels**. The diameter of the spheroids was quantified using ImageJ software on the day 7 of 3D cell culture showing cellular aggregates of variable size. The range of spheroids diameter was 100-400 µm. The data indicating that amyloid hydrogels mediated spheroids were larger than those formed in Matrigel for all the five cell types. n=4 independent experiments.

**Supplementary figure 9:**

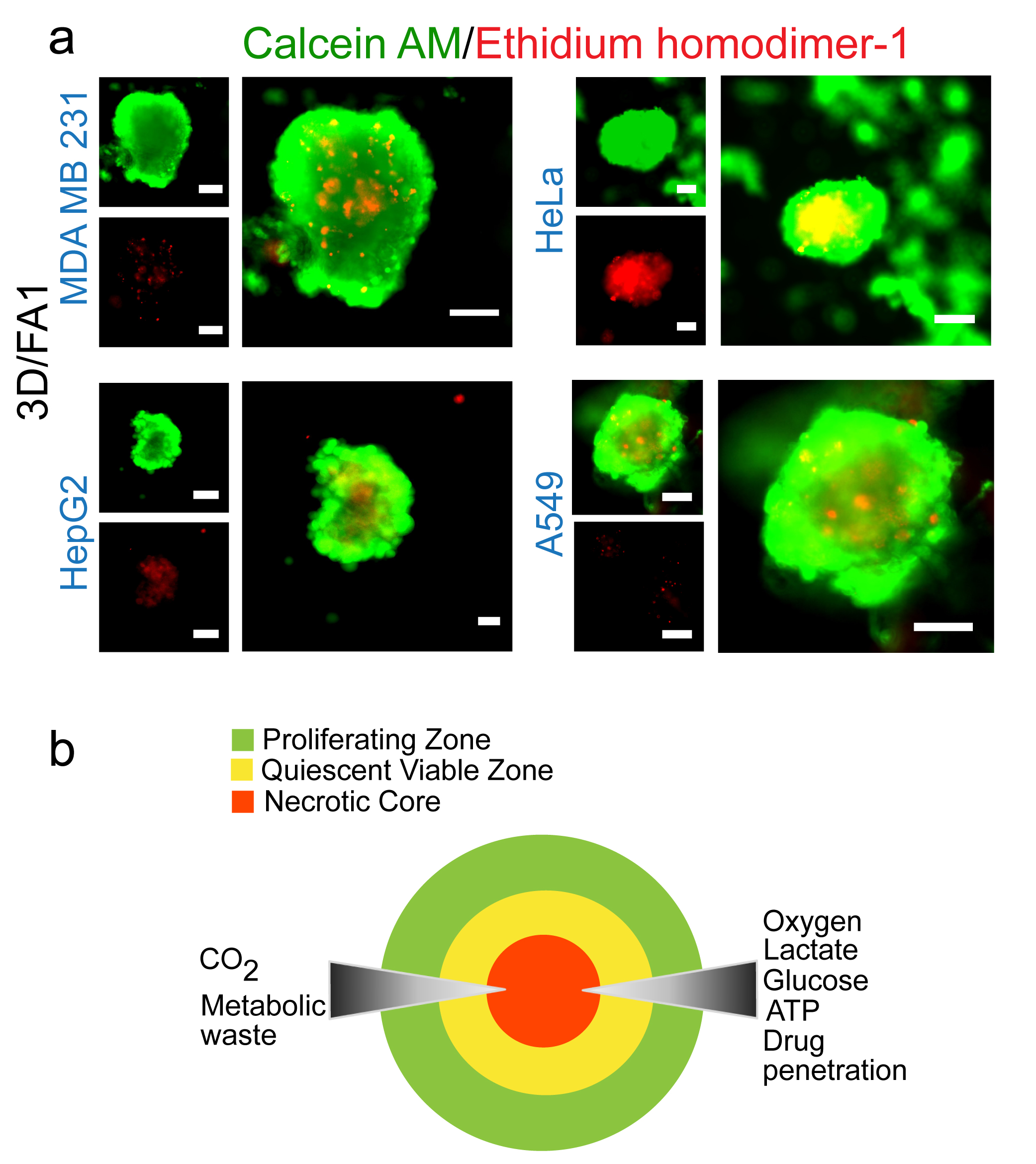

**Figure S9. Cell viability of cancer cells spheroids with the amyloid hydrogels.** (a) Calcein AM/Ethidium homodimer-1 staining (cell viability assay) of MDA MB 231, HeLa, HepG2, and A549 cancer cell aggregates on day 7 showing viable cells (green) and non-viable cells (red). Merged confocal image of the spheroids exhibiting spherical geometry with a concentric organization of proliferating, quiescent and dead cells. The scale bar is 100 µm. n=3 independent experiments. (b) Schematic representation of different zones of the spheroids (> 250 um in size), with densely packed cells at the center of the spheroid representing necrotic core, mid-region quiescent viable zone, and outermost proliferating region. Three independent experiments were performed.

**Supplementary figure 10:**

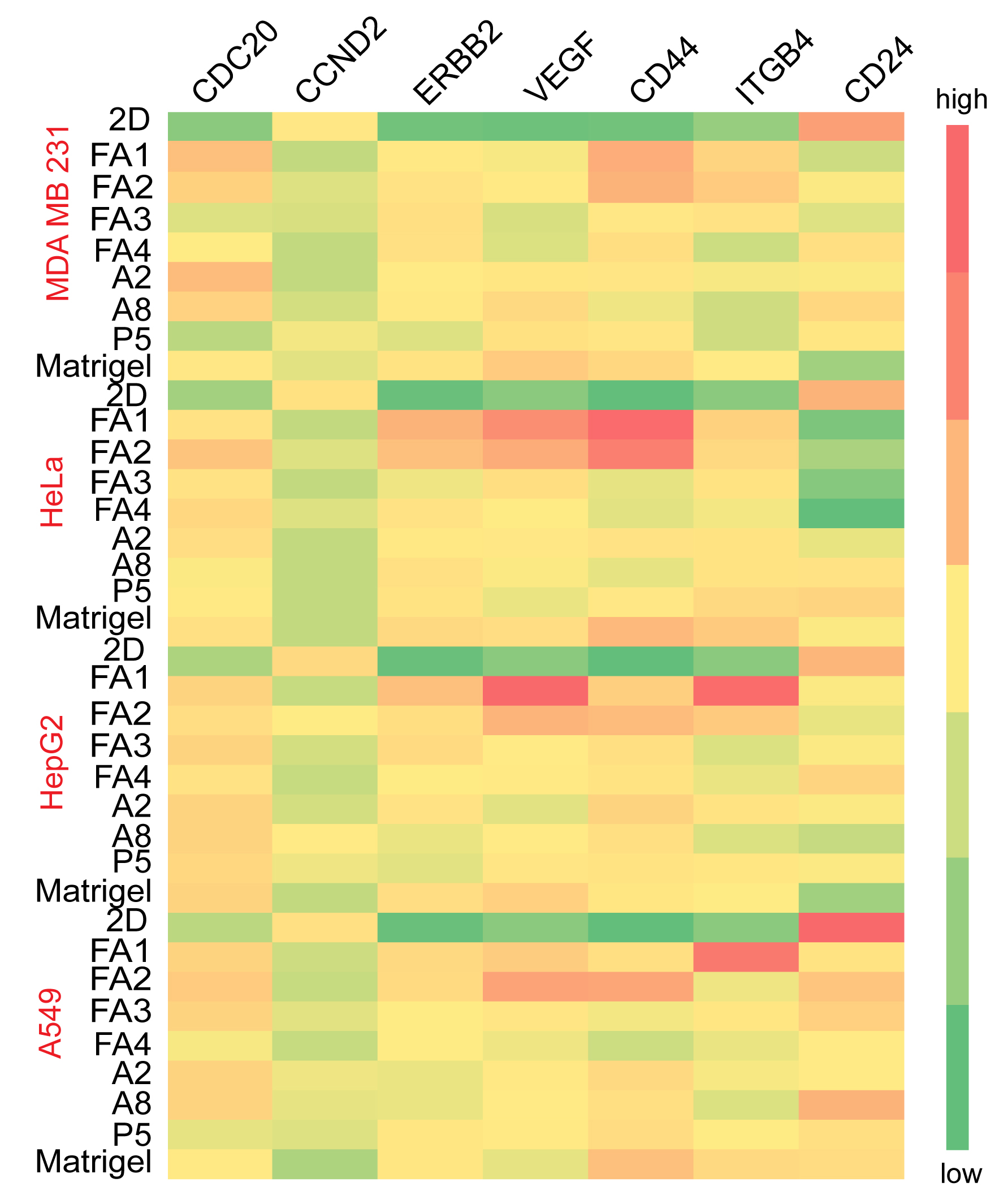

**Figure S10. Gene expression profile of MDA MB 231, HeLa, HepG2 and A549 cells in 3D cell culture with various amyloid hydrogels.** Heat map illustrating RT-qPCR data of potent cancer biomarkers expression in 3D cell culture in comparison to 2D cell culture. Matrigel was used as control. Red and green colours indicating upregulation and downregulation of gene expressions in cancer cells in compared to corresponding 2D monolayer cells. Data showing upregulation of pro-oncogenic markers such as ERBB2, VEGF, CD44, CDC 20, and ITGB4 along with downregulation of CD24 and CCND2. Three independent experiments were performed.

**Supplementary figure 11:**

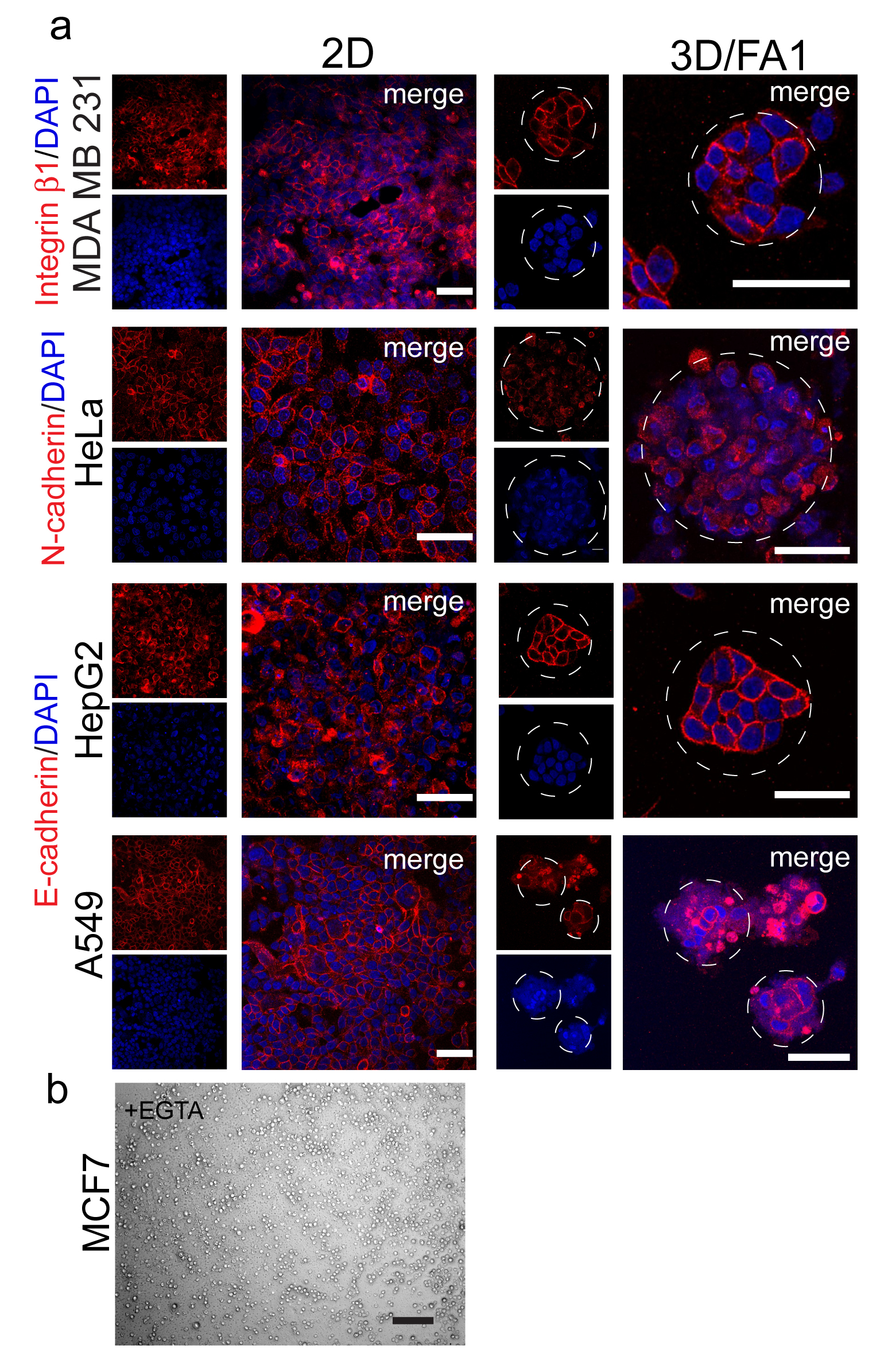

**Figure S11. Immunofluorescence staining of cell adhesion protein markers in 3D culture.** Immunofluorescence images showing expression of β1-integrin, E-cadherin, and N-cadherin in MDA MB 231, HepG2, A549, and HeLa cells, respectively, in 3D and 2D cell culture. The scale bar is 100 µm. Three independent experiments were performed. (b) Bright-field image of MCF7 cells cultured for 5 days with FA1 amyloid hydrogel in the presence of EGTA showing no cellular aggregation. The scale bar is 100 µm. n=3 independent experiments.

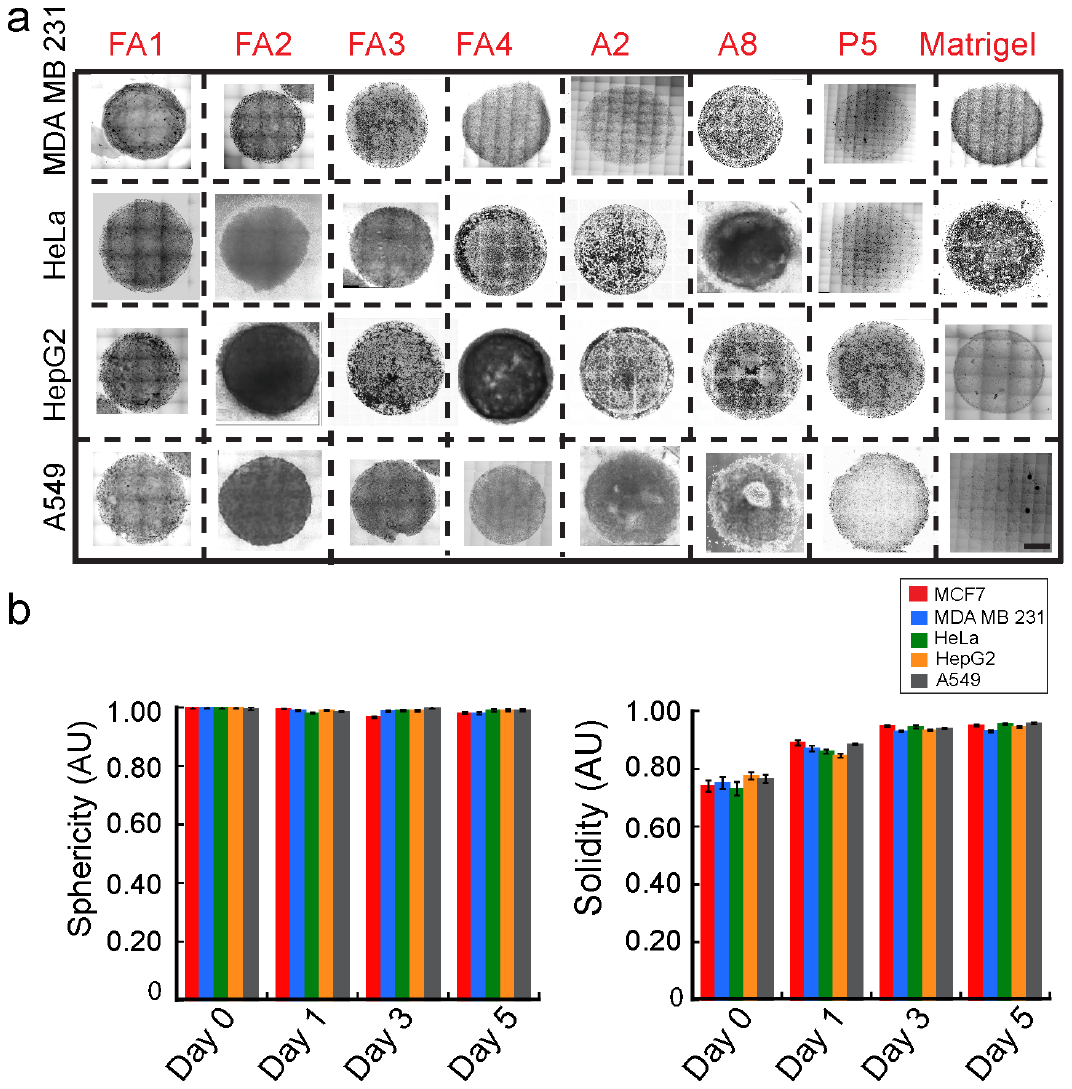

**Supplementary figure 12:**

**Figure S12. High-throughput spheroid formation for drug screening using drop cast method.** (a) Bright-field images showing spheroids formation by all five cancer cell lines with all amyloid hydrogels on day 3. Images showing the spherical and relatively similar size of spheroids in different gels. The scale bar is 500 µm. n=5 independent experiments. (b) Circularity (sphericity) of cellular spheroids with FA1 amyloid hydrogel was calculated from day 1 to day 5. Drop cast spheroids remain spherical throughout the culturing period. (c) Cellular density (solidity) analysis showing the degree of compaction by cells inside the drop cast spheroids for high-throughput drug assays. Data showed that the solidity increased from day 1 (0.7) to day 3 (0.95) and remains stable till day 5. The data plotted represent mean ± s.e.m, n=5 independent experiments.

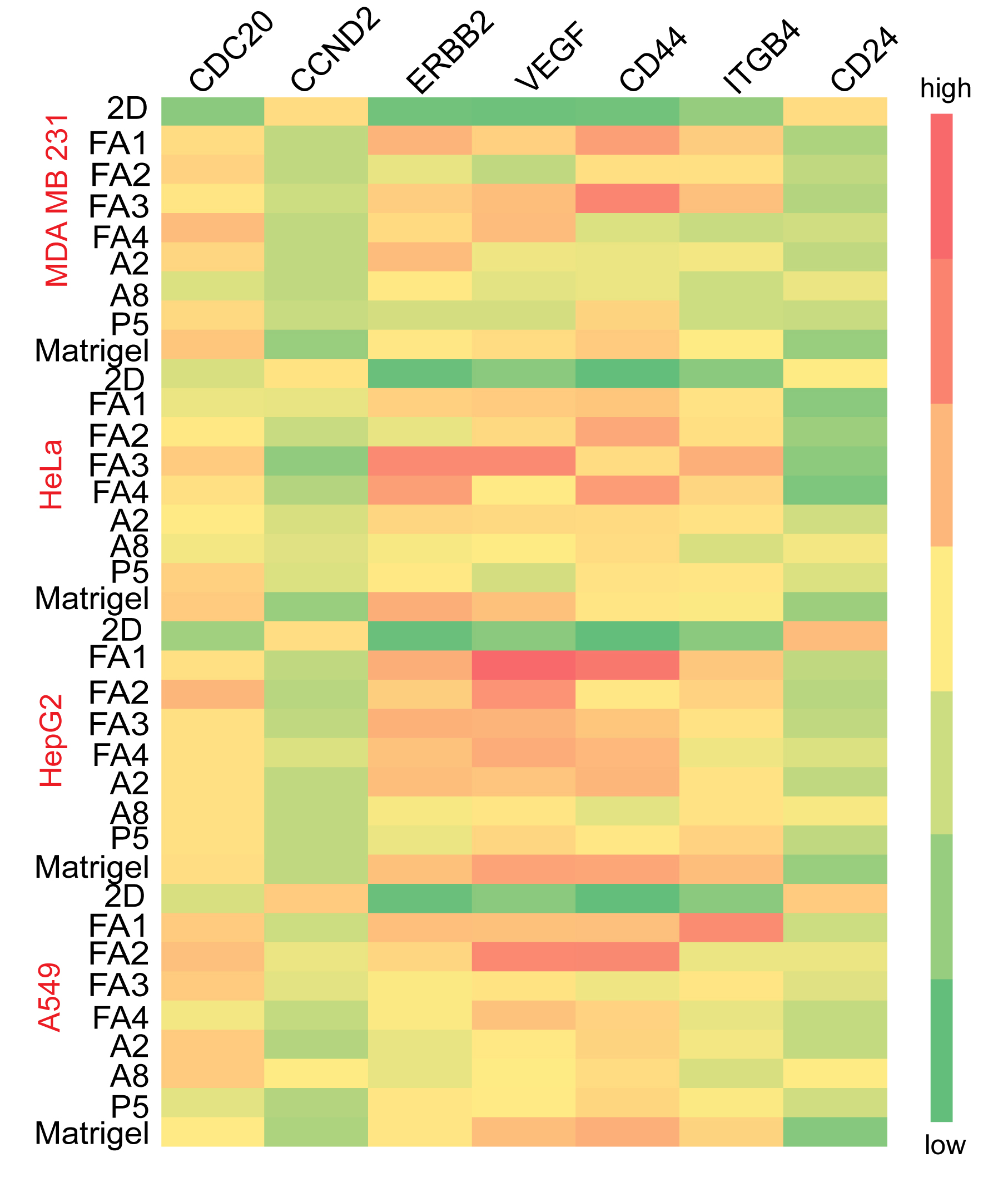
**Supplementary figure 13:**

**Figure S13. Gene expression profile of MDA MB 231, HeLa, Hep G2, and A549 spheroids cultured with amyloid hydrogels using the drop cast method.** Heat map illustrating RT-qPCR data of potent cancer biomarkers expression in 3D drop cast spheroid formed by cancer cells in comparison to 2D culture. Matrigel was used as control. Red and green colours indicating upregulation and downregulation of gene expressions in cancer cells in compared to corresponding 2D monolayer cells, respectively. Data showing up-regulation of pro-oncogenic markers such as ERBB2, VEGF, CD44, CDC 20, and ITGB4 along with downregulation of CD24 and CCND2. Three independent experiments were performed.

**Supplementary figure 14:**

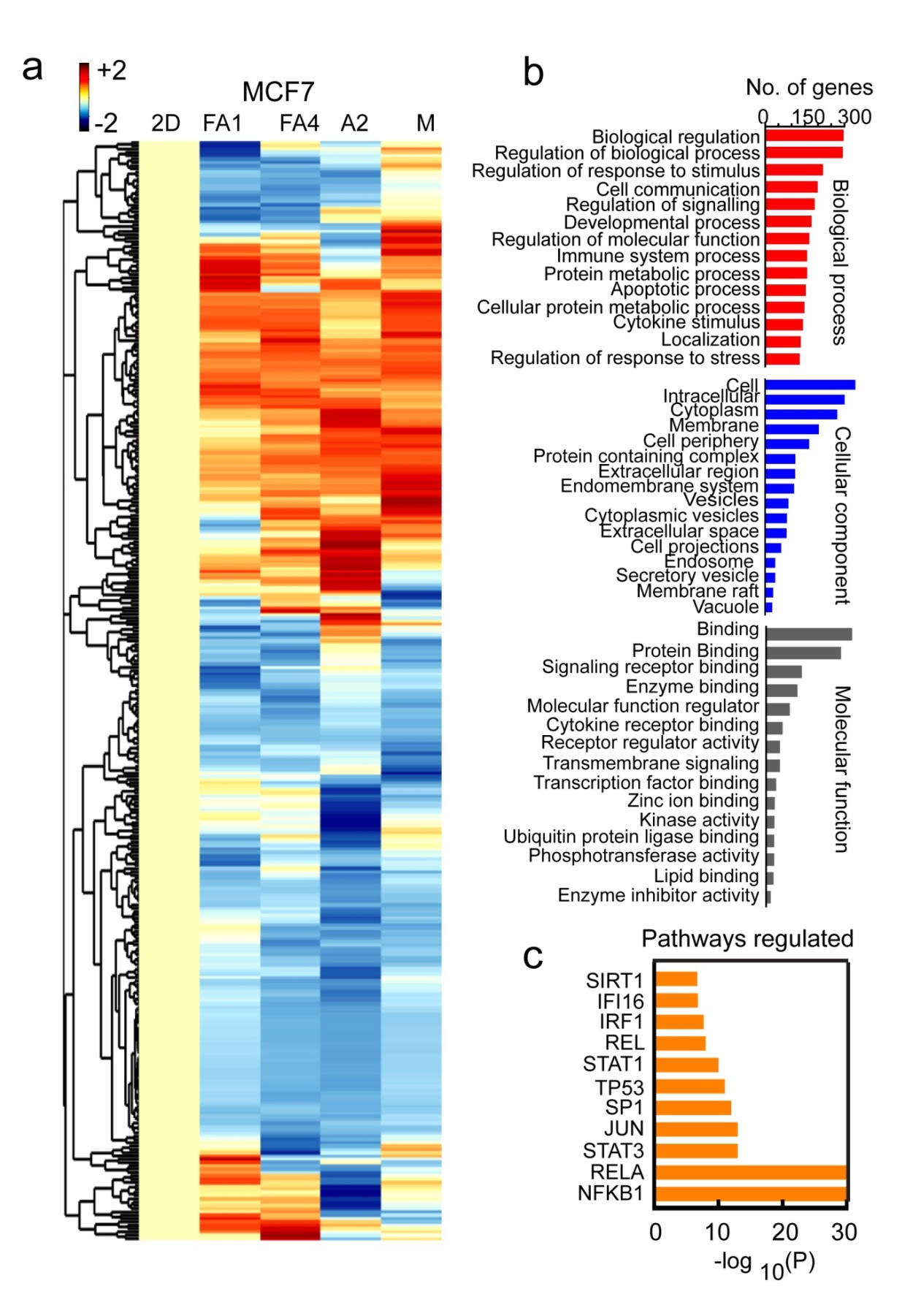

**Figure S14: Gene expression and pathway analysis of 3D MCF7 spheroid formed by drop cast method** (a) Heat map illustrating hierarchical clustering of breast cancer biomarker gene expressions array in 3D MCF7 spheroids (3D drop cast method) in comparison to 2D monolayer culture. Matrigel was used as control. Analysis of row dendrograms indicating that FA1 gel was the closest match with Matrigel compared to others and thereby clustered together. Red and blue indicating higher and lower expression of the genes, respectively in MCF7 spheroid in comparison to 2D monolayer culture. The intensity of the colours is proportional to a log2 of the fold changes. (b) Gene ontology study showing differentially regulated genes responsible for different biological processes, cellular components, and molecular functions. (c) Pathway enrichment analysis showing various signaling pathways (such as NFkB-1, STAT3, JUN, RELA) functioning in spheroid aggregation.

**Supplementary figure 15:**

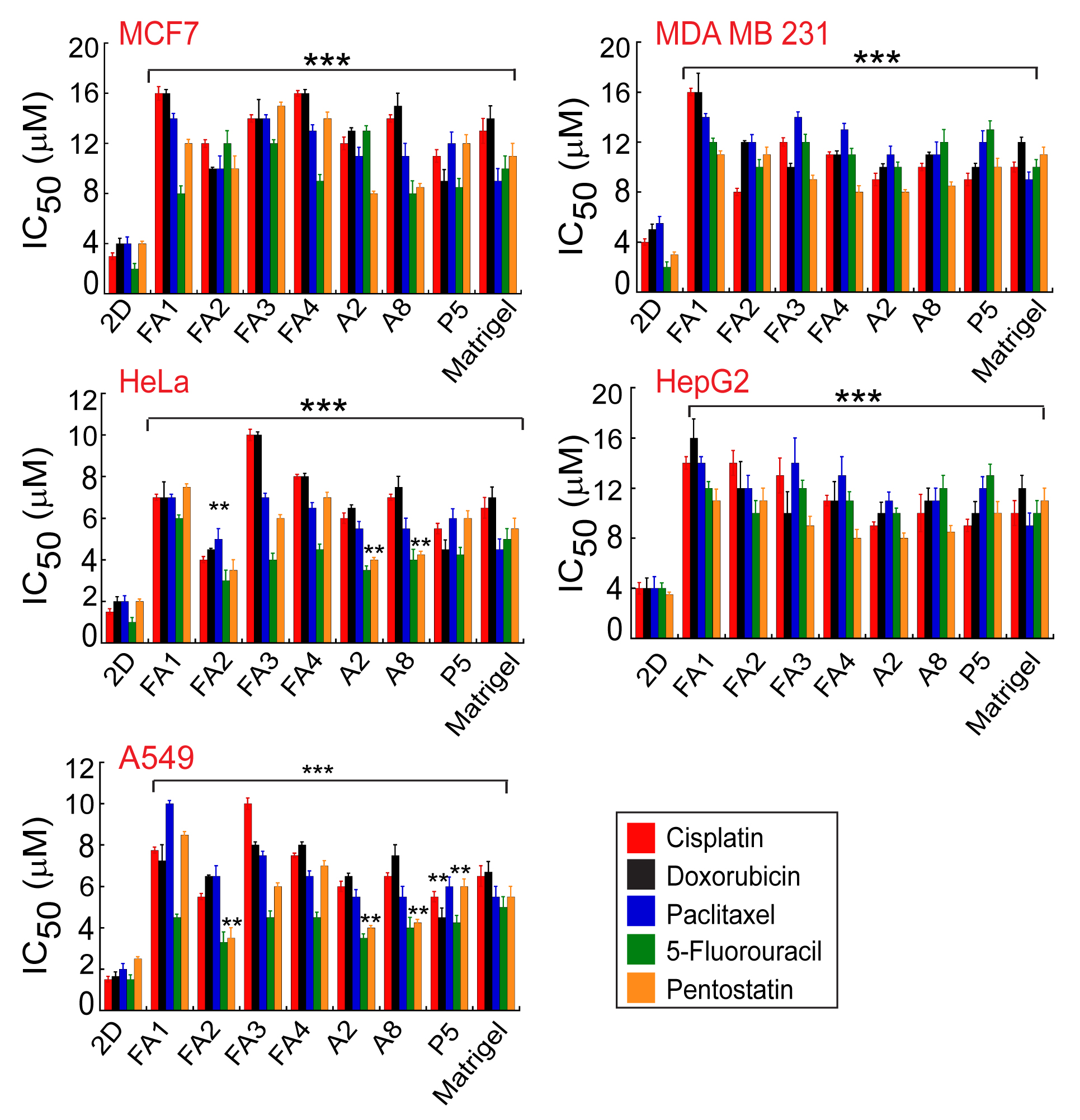

**Figure S15. Cell toxicity of spheroid by five different chemotherapeutic drugs using MTT assay**. The bar diagram (IC_50_) showing many folds higher cell viability in presence of different doses of various chemotherapeutic drugs after 24 h of treatment. The plot of IC_50_ of the anti-cancer drug for 2D was approximately 3 to 4-fold lower indicating that spheroids were more resistant to drugs as compared to 2D monolayer cells. The values plotted represent mean ± s.e.m, n=4 independent experiments. Statistical significance (***p ≤ 0.001, **p ≤ 0.01) is determined by one-way ANOVA followed by Dunnett's multiple comparison post-hoc test.

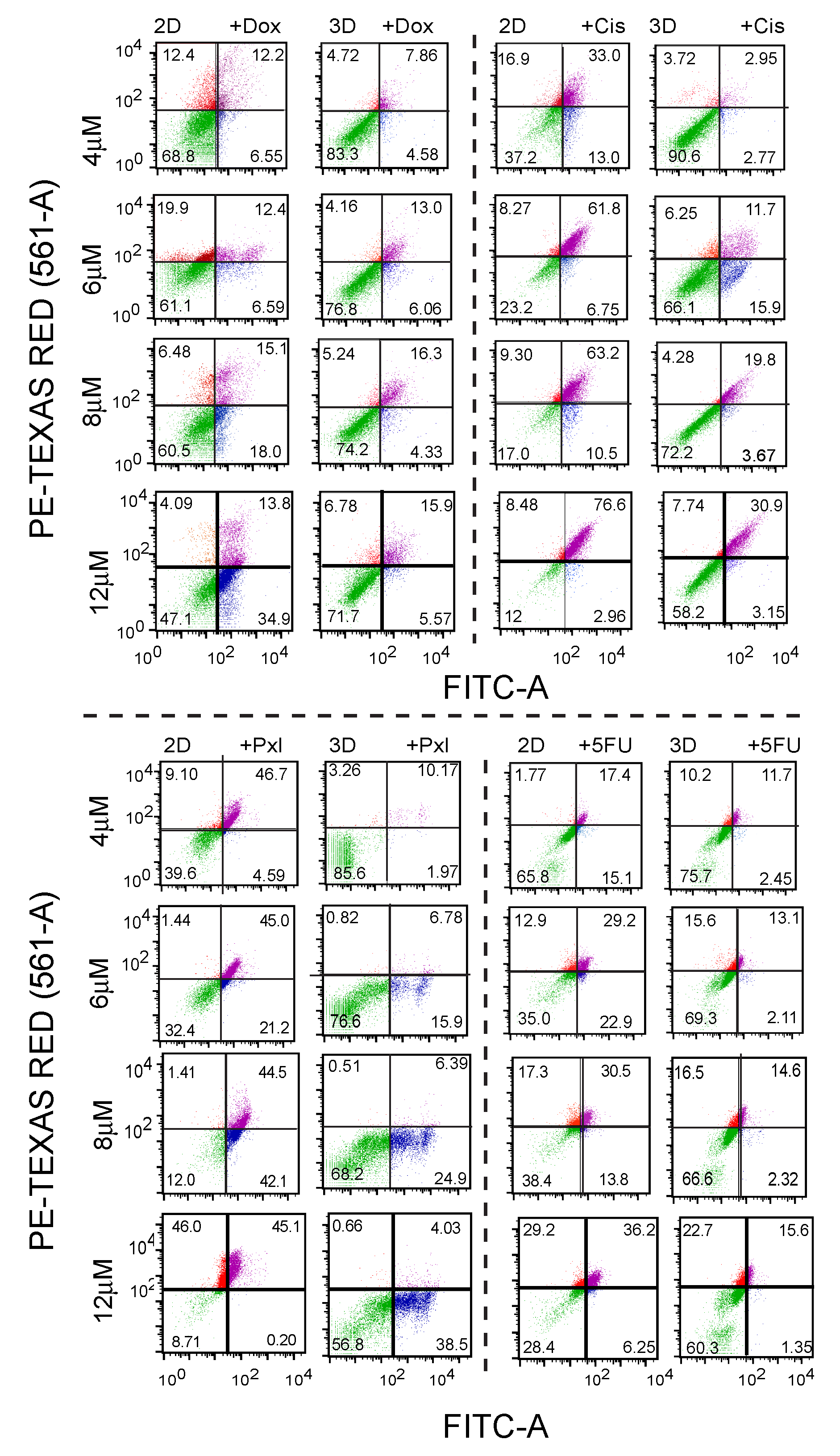
**Supplementary figure 16:**

**Figure S16. FACS analysis of cellular toxicity in presence of various chemotherapeutic drugs.** Scatter plots of FACS analysis with Annexin V/PI showing cell death in 2D monolayer and 3D MCF7 spheroids cultured with FA1 hydrogel in presence of doxorubicin, cisplatin, paclitaxel, and 5-Fluorouracil (4, 6 and 12 µM) for 24 h. The data showing higher cell death of 2D cultured cells in comparison to a 3D spheroid. The data suggesting that dense spheroids formed in 3D cell culture prevented apoptosis or death when treated with the anti-cancer drugs. Cells were classified into four groups based on dye uptake: viable cells (lower left quadrant), early apoptotic cells (lower right quadrant), late apoptotic cells (upper right quadrant), and necrotic cells (upper left quadrant). Three independent experiments were performed.

**Supplementary figure 17:**

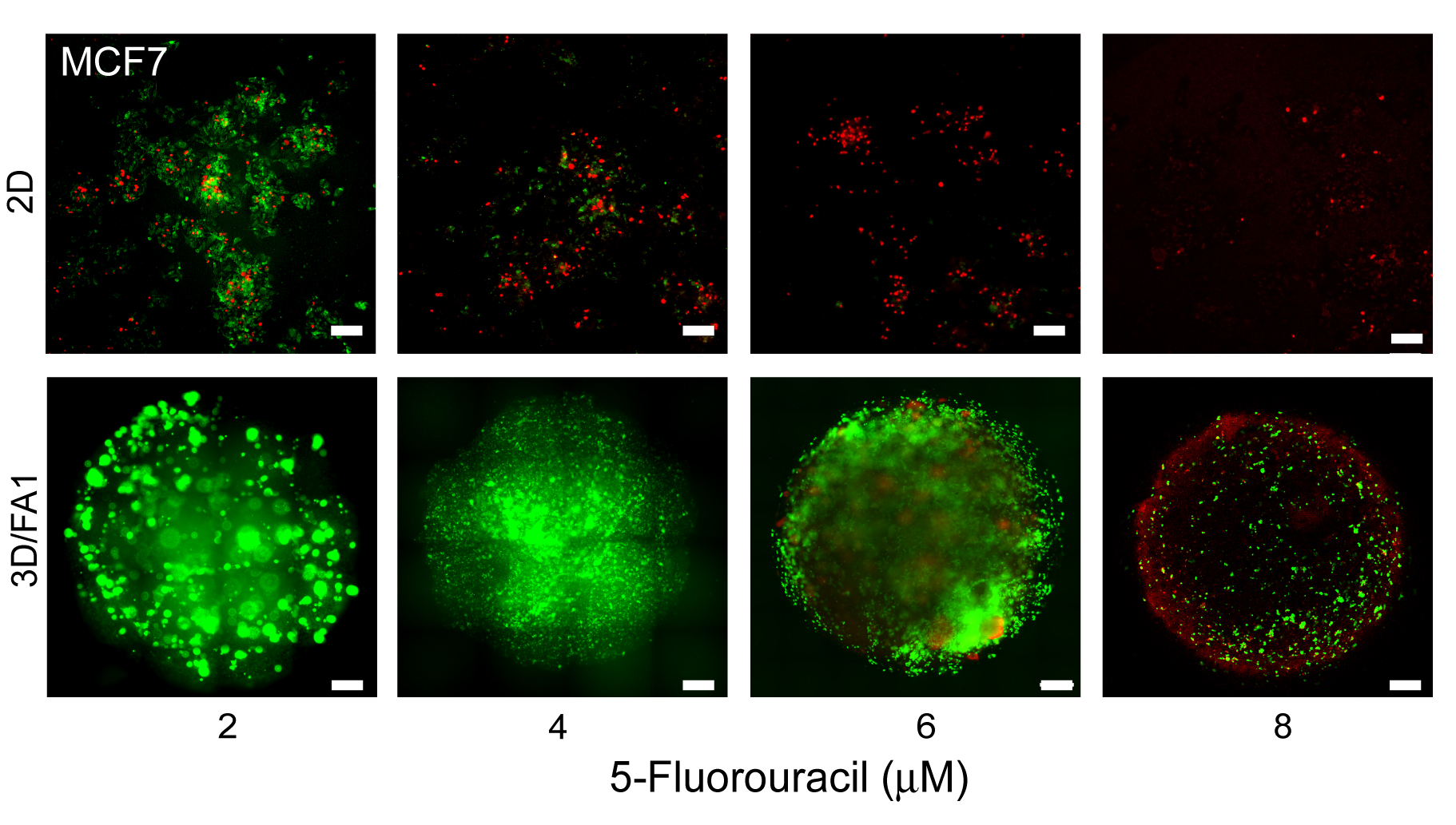

**Figure S17. Fluorescent images of MCF7 cell spheroids treated with different concentrations of 5-fluorouracil.** Cellular toxicity (MCF7 cells) using Calcein AM/Ethidium homodimer-1 followed by fluorescence microscopy showing drop cast spheroids were resistant to cell death in comparison to 2D monolayer culture. Live cells were green (Calcein AM), while dead cells were stained red (Ethidium homodimer-1). The data indicating the compact structure of the 3D spheroid not allowing the drug to penetrate and cell death is restricted to the outer cell layers only. n=3 independent experiments.

**Supplementary figure 18:**

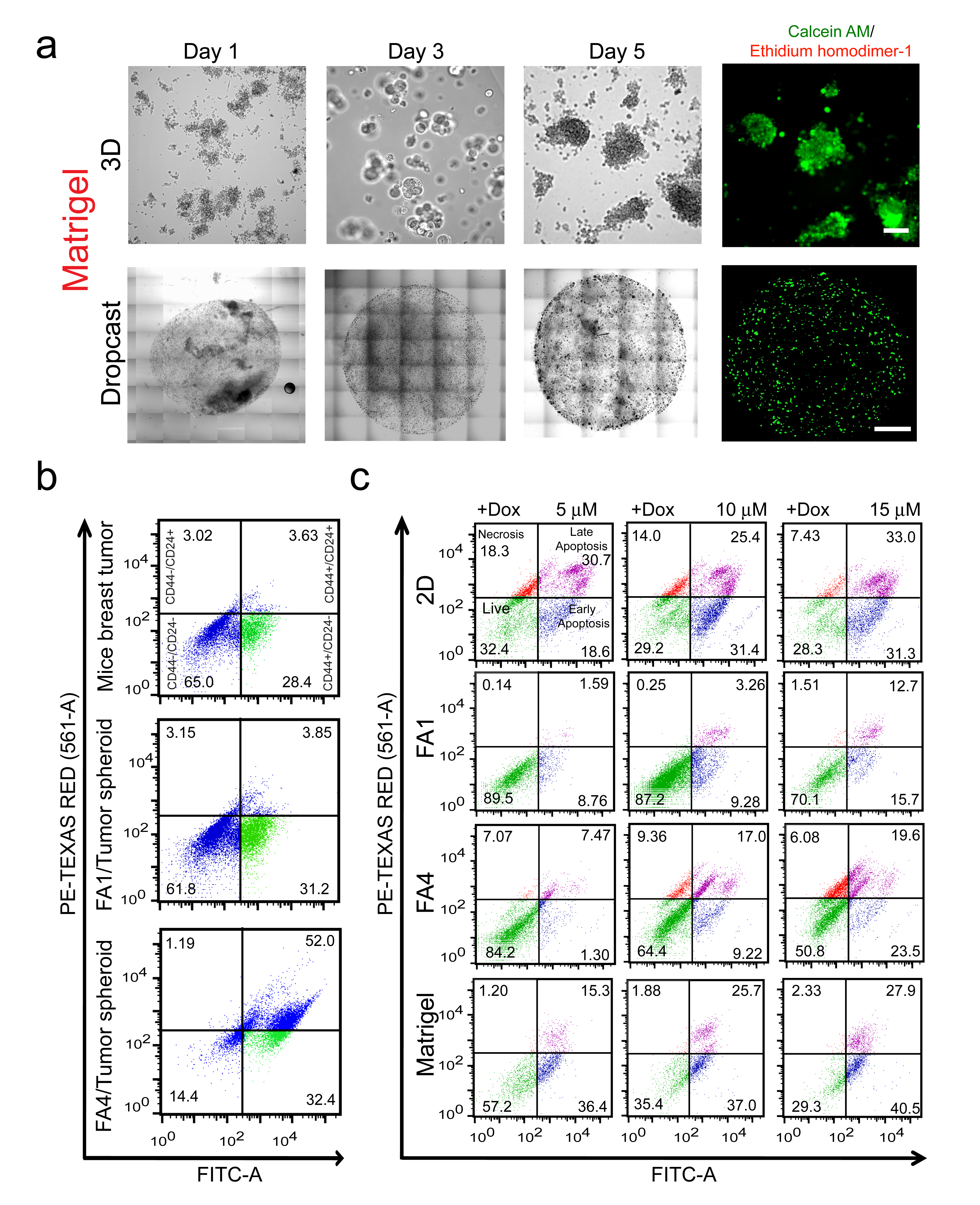

**Figure S18. Tumor spheroids from mice xenograft tumor.** (a) Time-dependent cellular aggregation in 3D culture (upper panel) and spheroid formation in drop cast method (lower panel) using Matrigel showing the growth of tumor spheroids by isolated cells from mice xenograft breast tumor. Corresponding fluorescence images of cell viability of the day 5 culture were stained with Calcein AM/Ethidium homodimer-1 are shown. The scale bar for 3D culture is 500 µm and the drop cast culture is 100 µm. (b) FACS sorting of CD44^+^/CD24^-^ population from day 5 spheroid cells showing similar CD44^+^/CD24^-^ cell population in compared with cells isolated from the original tumor. (b) Cell death assay in the presence of doxorubicin using Annexin V/PI followed by FACS analysis. Data showing differential cell death in tumor spheroids with FA1 hydrogel versus 2D monolayer culture, suggesting densely packed spheroids formed in 3D cell culture are more resistant to apoptosis/death. Scattering of cells within the quadrant is due to the presence of different sizes of cells that are isolated from tumors. Three independent experiments were performed.

**Supplementary tables**

**Supplementary Table S1. List of primers used for real-time qPCR for gene expression study for 3D spheroids compared to the 2D monolayer**

| **Gene** | **Function** | **Sequences of primers** |
| --- | --- | --- |
| GAPDH | Housekeeping ^17^ | Forward: 5’-CATTTTACGCTGATCCAGG-3’  Reverse: 5’-GGGTTCGAAATGAGGATG-3’ |
| CD24 | Cell adhesion, tumor suppressor genes ^18^ | Forward: 5′-TGCTCCTACCCACGCAGATT-3’  Reverse: 5’- GGCCAA CCCAGAGTTGGAA-3′ |
| CD44 | Cell–cell interactions, cell adhesion and migration, cancer stem cells ^18^ | Forward: 5’- GATCATCTTGGCATCCCTCT-3’  Reverse: 5’- TGAGTCCACTTGGCTTTCTG-3’ |
| CDC20 | Cell division regulator ^19^ | Forward: 5’-GAGCTTTGGAATCTCAGAATG-3’  Reverse: 5’-CAGGTCCCAAATTTTCACTG-3’ |
| VEGF | Angiogenesis, epithelial-mesenchymal transition ^20^ | Forward: 5’- GCAGCTTGAGTTAAACGAACG-3’  Reverse: 5’- GGTTCCCGAAACCCTGAG-3’ |
| CCND2 | Cell cycle regulation, differentiation, tumor suppressor genes^21^ | Forward: 5’-ACTTCATTGAGCACATCTTG-3’  Reverse: 5’-ACATGGCAAACTTAAAGTCG-3’ |
| ERBB2 | Proto-oncogene, cell growth, and survival^12^ | Forward: 5’- ACCTGCTGAACTGGTGTATG-3’  Reverse: 5’-TGACATGGTTGGGACTCTTG-3’ |
| ITGB4 | Cell extracellular matrix interactions, cell growth, migration, and/or apoptosis^22^ | Forward: 5’-ACTACACCCTCACTGCAGAC-3’  Reverse: 5’-TCTGGCTTGCTCCTTGATGA-3’ |
| CDH1 | Cell-cell interactions, tumor progression, and metastasis ^23^ | Forward: 5’-GAACGATTGCCACATACAC-3’  Reverse: 5’-GAATTCGGGCTTGTTGTCAT-3’ |
| SLUG | Epithelial-mesenchymal transition, drug resistance^24^ | Forward: 5’-TGTTTGCAAGATCTGCGGC-3’  Reverse: 5’-TGCAGTCAGGGCAAGAAAAA-3’ |
| CTNNB1 | Proto-oncogene, tumor progression ^25^ | Forward: 5’-GTTCGCCTTCACTATGGACTACC-3’  Reverse: 5’-GGACCCCTGCAGCTACTCTTT -3’ |

**Supplementary Table S2: Differential expression of genes in spheroids formed with amyloid hydrogels and Matrigel.**

| **Sample** | **Upregulated genes** | **Downregulated genes** |
| --- | --- | --- |
| FA1 | 3303 | 1937 |
| FA4 | 3386 | 2040 |
| A2 | 3401 | 4157 |
| Matrigel | 3440 | 1751 |

**Supplementary Table S3: Breast cancer organoid media recipe.**

The tumor spheroids were grown in organoid media recipe mentioned below^16^.

| **Medium component** | **Final concentration** |
| --- | --- |
| R-Spondin 3 | 250 ng·ml^-1^ |
| Neuregulin 1 | 5 nM |
| FGF 7 | 5 ng·ml^-1^ |
| FGF 10 | 20 ng·ml^-1^ |
| EGF | 5 ng·ml^-1a^ |
| Noggin | 100 ng·ml^-1^ |
| A83-01 | 500 nM |
| Y-27632 | 5 mM |
| SB202190 | 500 nM^b^ |
| B27 supplement | 1x |
| N-Acetylcysteine | 1.25 mM |
| Nicotinamide | 5 mM |
| GlutaMax 100x | 1x |
| Hepes | 10 mM |
| Penicillin/Streptomycin | 100 U·ml^-1^ / 100 mg·ml^-1^ |
| Primocin | 50 µg·ml^-1^ |
| Advanced DMEM/F12 | 1x |

**References**

1.	Das, S. et al. Implantable amyloid hydrogels for promoting stem cell differentiation to neurons. *NPG Asia Mater.* **8**, e304-e304 (2016).

2.	Das, S., Kumar, R., Jha, N.N. & Maji, S.K. Controlled Exposure of Bioactive Growth Factor in 3D Amyloid Hydrogel for Stem Cells Differentiation. *Adv. Healthc. Mater.* **6** (2017).

3.	Jacob, R.S. et al. Self healing hydrogels composed of amyloid nano fibrils for cell culture and stem cell differentiation. *Biomaterials* **54**, 97-105 (2015).

4.	Maji, S.K. et al. Functional amyloids as natural storage of peptide hormones in pituitary secretory granules. *Science* **325**, 328-332 (2009).

5.	Wei, G. et al. Self-assembling peptide and protein amyloids: from structure to tailored function in nanotechnology. *Chem. Soc. Rev.* **46**, 4661-4708 (2017).

6.	Fernandez-Escamilla, A.M., Rousseau, F., Schymkowitz, J. & Serrano, L. Prediction of sequence-dependent and mutational effects on the aggregation of peptides and proteins. *Nat. Biotechnol.* **22**, 1302-1306 (2004).

7.	Fields, G.B. & Noble, R.L. Solid phase peptide synthesis utilizing 9-fluorenylmethoxycarbonyl amino acids. *Int. J. Pept. Protein. Res.* **35**, 161-214 (1990).

8.	Liang, C.-C., Park, A.Y. & Guan, J.-L. In vitro scratch assay: a convenient and inexpensive method for analysis of cell migration in vitro. *Nat. Protoc.* **2**, 329-333 (2007).

9.	Lin, R.-Z. & Chang, H.-Y. Recent advances in three-dimensional multicellular spheroid culture for biomedical research. *Biotechnol. J.* **3**, 1172-1184 (2008).

10.	Eilenberger, C., Rothbauer, M., Ehmoser, E.-K., Ertl, P. & Küpcü, S. Effect of Spheroidal Age on Sorafenib Diffusivity and Toxicity in a 3D HepG2 Spheroid Model. *Sci. Rep.* **9**, 4863 (2019).

11.	Sirenko, O., Hesley, J., Rusyn, I. & Cromwell, E.F. High-content high-throughput assays for characterizing the viability and morphology of human iPSC-derived neuronal cultures. *Assay Drug Dev. Technol.* **12**, 536-547 (2014).

12.	Slamon, D.J. et al. Studies of the HER-2/neu proto-oncogene in human breast and ovarian cancer. *Science* **244**, 707-712 (1989).

13.	Kim, H., Phung, Y. & Ho, M. Changes in Global Gene Expression Associated with 3D Structure of Tumors: An Ex Vivo Matrix-Free Mesothelioma Spheroid Model. PloS one **7**, e39556 (2012).

14.	Huang, D.W. et al. The DAVID Gene Functional Classification Tool: a novel biological module-centric algorithm to functionally analyze large gene lists. *Genome Biol.* **8**, R183-R183 (2007).

15.	Zhou, Y. et al. Metascape provides a biologist-oriented resource for the analysis of systems-level datasets. *Nat. Commun.* **10**, 1523 (2019).

16.	Sachs, N. et al. A Living Biobank of Breast Cancer Organoids Captures Disease Heterogeneity. *Cell* **172**, 373-386.e310 (2018).

17.	Barber, R.D., Harmer, D.W., Coleman, R.A. & Clark, B.J. GAPDH as a housekeeping gene: analysis of GAPDH mRNA expression in a panel of 72 human tissues. *Physiol. Genomics* **21**, 389-395 (2005).

18.	Chen, C., Zhao, S., Karnad, A. & Freeman, J.W. The biology and role of CD44 in cancer progression: therapeutic implications. *J. Hematol. Oncol.* **11**, 64-64 (2018).

19.	Kidokoro, T. et al. CDC20, a potential cancer therapeutic target, is negatively regulated by p53. *Oncogene* **27**, 1562-1571 (2008).

20.	Carmeliet, P. VEGF as a key mediator of angiogenesis in cancer. *Oncology* **69** 3, 4-10 (2005).

21.	McCullough, L.E. et al. Modification of the association between recreational physical activity and survival after breast cancer by promoter methylation in breast cancer-related genes. *Breast Cancer Res.* **19**, 19 (2017).

22.	Chung, J., Bachelder, R.E., Lipscomb, E.A., Shaw, L.M. & Mercurio, A.M. Integrin (alpha 6 beta 4) regulation of eIF-4E activity and VEGF translation: a survival mechanism for carcinoma cells. *J. Cell Biol.* **158**, 165-174 (2002).

23.	Jeanes, A., Gottardi, C.J. & Yap, A.S. Cadherins and cancer: how does cadherin dysfunction promote tumor progression? *Oncogene* **27**, 6920-6929 (2008).

24.	Pérez-Mancera, P.A. et al. SLUG in cancer development. *Oncogene* **24**, 3073-3082 (2005).

25.	Zhan, T., Rindtorff, N. & Boutros, M. Wnt signaling in cancer. *Oncogene* **36**, 1461-1473 (2017).
